# An environmentally ultrasensitive fluorine probe to resolve protein conformational ensembles by ^19^F NMR and cryo-EM

**DOI:** 10.1101/2022.03.29.486269

**Authors:** Yun Huang, Krishna Reddy, Clay Bracken, Biao Qiu, Wenhu Zhan, David Eliezer, Olga Boudker

## Abstract

Limited chemical shift dispersion represents a significant barrier to studying multi-state equilibria of large membrane proteins by ^19^F NMR. We describe a novel monofluoroethyl ^19^F probe that dramatically increases the chemical shift dispersion. The improved conformational sensitivity and line shape enable the detection of previously unresolved states in 1D NMR spectra of a 134 kDa membrane transporter. Changes in the populations of these states in response to ligand binding, mutations, and temperature correlate with population changes of distinct conformations in structural ensembles determined by single-particle cryo-electron microscopy. Thus, ^19^F NMR can guide sample preparation to discover and visualize novel conformational states and facilitate image analysis and 3D classification.

## Introduction

The functions of numerous membrane proteins, including transporters, channels, and receptors, require conformational transitions. Insights into the structure and dynamics of the functional states are crucial for mechanistic understanding and therapeutic development ^1–3^. Crystallography and single-particle cryogenic electron microscopy (cryo-EM) can provide structural snapshots of different states in a protein ensemble, while various bulk ^4^ and single-molecule ^5–7^ approaches can report on dynamics. Nuclear magnetic resonance (NMR) spectroscopy is a powerful tool for studying the dynamics of small proteins. However, NMR sensitivity and resolution decline with protein size, and probing the dynamics of proteins over 100 kDa, including membrane proteins, by NMR remains challenging.

1D ^19^F NMR using site-specific probes can be effective for this purpose because of its high sensitivity, the absence of background signals, the exquisite responsiveness of ^19^F chemical shifts to conformational changes, and its compatibility with inexpensive low-field instruments ^8–10^. Furthermore, 1D ^19^F NMR is a single-pulse experiment with no heteronuclear magnetization transfer. It can therefore quantify state populations of even weak or broad resonances. The unique advantages of ^19^F NMR have led to wide-ranging applications in studying membrane protein dynamics ^9, 11^, protein aggregation ^12^, protein behavior in cells ^13^, and drug screening ^14^. Commonly used ^19^F probes include aromatic fluorine and trifluoromethyl groups. The latter show faster longitudinal and slower transverse relaxation rates due to fast rotation, leading to higher sensitivity ^15^. Manglik et al. developed a trifluoromethyl phenyl group, where the phenyl ring functions as a chemical shift dispersion amplifier ^16–17^. However, the chemical shift dispersion of trifluoromethyl-based probes generally does not exceed 2 ppm ^16, 18–20^, resulting in severe resonance overlap in large proteins. Recently, Boeszoermenyi et al. reported ^19^F-^13^C TROSY, improving resonance dispersion via a second dimension ^21^, but sacrificing the advantages of the 1D experiment. Therefore, developing ^19^F probes with improved dispersion, i.e., increased environmental sensitivity, remains a challenge and is essential for expanding ^19^F NMR applications ^17, 22–23^.

Herein, we report a novel cysteine-conjugated ^19^F monofluoroethyl (mFE) probe exhibiting narrow linewidths and ultrahigh sensitivity to conformational changes with chemical shift dispersion reaching 9 ppm.

^19^F chemical shifts of aliphatic monofluorides depend on C-C-C-F dihedral angles in addition to local electric fields and van der Waals interactions ^24 25^. In a trifluoroalkyl probe, all rotamers are equivalent, but in the monofluoroalkyl probe, the *gauche*- and *anti*-rotamers, which can exhibit chemical shift differences of up to 14 ppm ^25^ are not equivalent. Because of the low energy barrier separating the rotamers, they exchange much faster than the NMR time scale ^26^, giving a population-weighted chemical shift. We hypothesized that for an mFE probe, weak ^19^F interactions with the local environment of distinct protein conformations might affect the equilibrium of the *gauche* and *anti*-isomers, giving rise to different population-weighted chemical shifts and larger chemical shift dispersions than for a trifluoroethyl (tFE) probe. Such weak interactions should not significantly alter the kinetics of rotamer exchange, which would remain fast on the NMR time scale. Additionally, fluorine chemical shift anisotropy (CSA) is approximately two times lower in mFE than in the tFE group ^27^. Because ^19^F transverse relaxation depends quadratically on the CSA, mFE might feature slower relaxation and narrower linewidths.

To test the environmental sensitivity of the novel mFE probe, we used the aspartate transporter Glt_Ph_, an excellent model protein because it samples several functional conformations with known structures. Glt_Ph_ is similar in size to or larger than many membrane receptors, channels, and transporters. As an archaeal homolog of human glutamate transporters, it harnesses energy from sodium gradients to transport aspartate via an “elevator” mechanism ^28^. It is an obligate homotrimer of 134 kDa molecular weight that can be reconstituted as a protein-micelle particle of ~300 kDa ^29^. During transport, the three Glt_Ph_ protomers function independently ^30^, undergoing conformational transitions between the outward- and inward-facing states (OFS and IFS), where the substrate-binding site can open to the extracellular solution and cytoplasm, respectively. ^19^F NMR measurements using mFE-labeled Glt_Ph_ resolve an unexpectedly complex conformational landscape, demonstrating the power of the new probe and prompting concordant cryo-EM studies that confirm multiple coexisting states. The response of the dominant NMR signals to environmental changes, including temperature, correlates with changes in the ensemble of structures determined by cryo-EM, allowing for provisional assignments of some of the NMR signals to specific structural states. Importantly, our data also suggest that cryo-EM can accurately reflect the effects of temperature and other conditions on conformational equilibria in solution, a subject of considerable controversy.

## Results

### High environmental sensitivity of the mFE probe

We designed and synthesized deuterated ^19^F labeling compounds S-(2-fluoroethyl-1,1,2,2-D_4_) and *p*-toluenethiosulfonate (TsSCD_2_CD_2_F) using commercially available reagents (**Fig. 1a** and **b**). Deuterated probes provide sharper lines resulting from reduced ^19^F-^2^H scalar couplings compared to ^19^F-^1^H couplings and from the elimination of ^19^F-^1^H dipole relaxation at the cost of reduced sensitivity due to longer longitudinal relaxation times (T_1_) and relaxation delays. We then prepared single cysteine mutants, A380C, A381C, and M385C, of a previously characterized fully functional cysteine-free Glt_Ph_ variant (termed wild-type, WT, for brevity; see Methods for details and Ref. ^20^) and a variant with additional gain-of-function mutations R276S/M395R (termed RSMR). The RSMR mutations accelerate transitions between OFS and IFS ^20, 31^ and substrate uptake by ∽8-fold ^32^. Thiosulfonates are known to react with cysteine thiols selectively, rapidly, and quantitatively (**Fig. 1c**) ^33^, and the observed efficiency of TsSCD_2_CD_2_F reacting with a single cysteine of Glt_Ph_ was near 100 % (**Fig. S1a**). All mFE-labeled mutants retained transport activity when reconstituted into liposomes (**Fig. S1b**).

**Figure 1.**
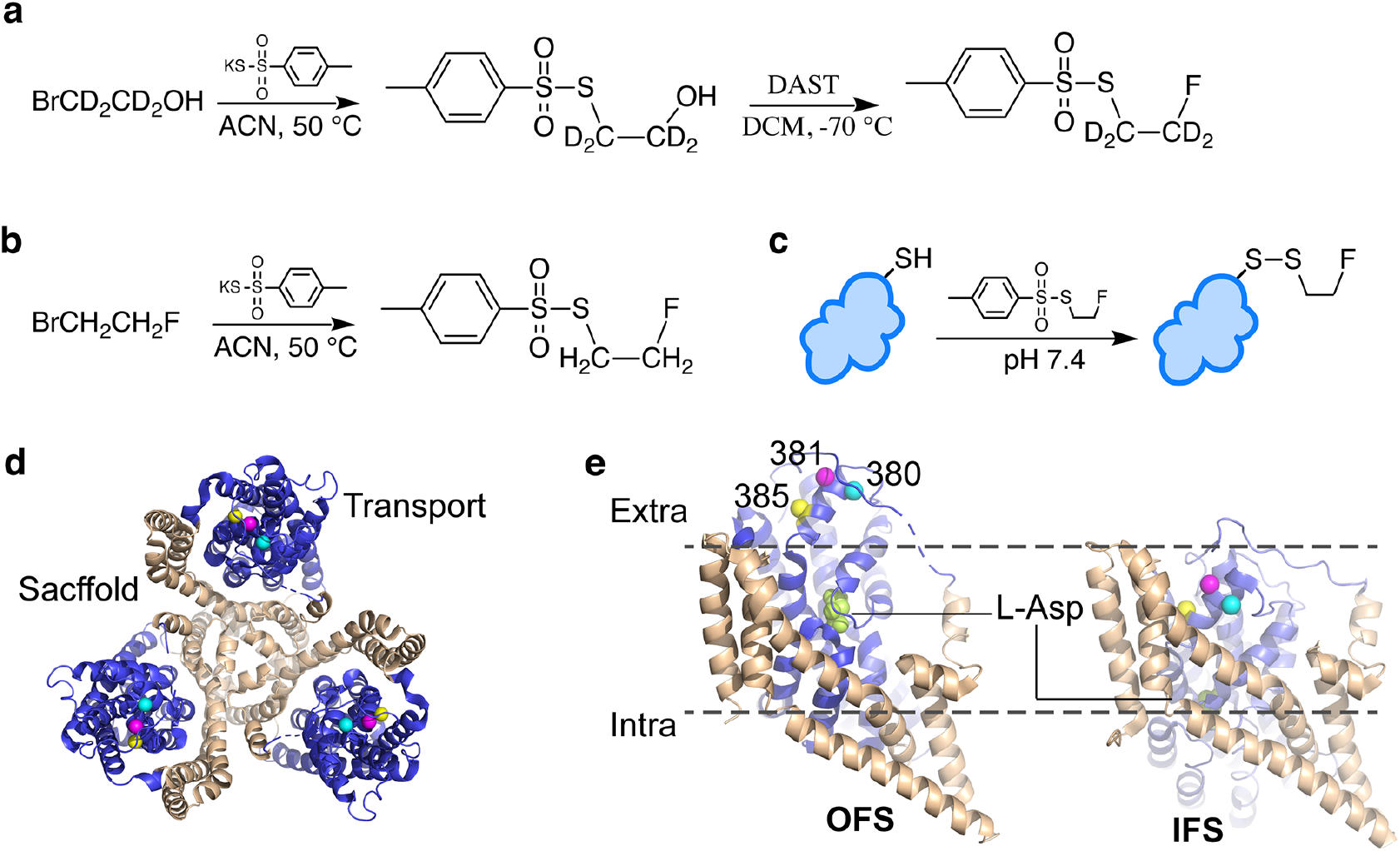
Synthesis of the mFE probes and site-specific protein labeling. Synthesis of ^19^F deuterated and protonated mFE probes TsSCD_2_CD_2_F (**a**) and TsSCH_2_CH_2_F (**b**). (**c**) Cysteine-specific labeling. (**d**) The structure of the Glt_Ph_ trimer with scaffold domains is colored tan, and the transport domains are blue. Colored spheres show the sites of single cysteine mutations A380C (cyan), A381C (magenta), and M385C (yellow). (**e**) The elevator transition of the Glt_Ph_ transport domain from OFS (left) to IFS (right). Single protomers are shown in the membrane plane, represented by the dotted lines. The bound substrate aspartate is shown as lime spheres. ACN: acetonitrile; DAST: diethylaminosulfur trifluoride; DCM: dichloromethane; X: hydrogen or deuterium; Ts: Toluenesulfonyl.

We next compared the chemical shift dispersion of the deuterated mFE and its predecessor tFE probe. We recorded 1D ^19^F spectra of mFE- and tFE-labeled WT-M385C and RSMR-M385C Glt_Ph_ in the presence of Na^+^ ions and aspartate or a competitive blocker (3S)-3-[[3-[[4-(trifluoromethyl)benzoyl]amino]phenyl]methoxy]-L-aspartic acid (TFB-TBOA). As previously reported, the tFE probe exhibited three peaks with a chemical shift dispersion of 1.1 ppm (**Fig. 2a** and **S2a**) ^20^. In contrast, the mFE spectra collectively featured five peaks, S1-S5, at 25 °C, with respective chemical shifts of ~ −214.8, 216.0, 216.6, 217.5, and 218.1 ppm (**Fig. 2b** and **S2b, c**). An analysis of three independently prepared samples showed that peak identification is highly reproducible, with only small deviations in the fitted peak positions, linewidths, and populations (**Fig. S3**). We ascribe the slight shift in the S4 peak in the presence of TFB-TBOA to a structural difference between the aspartate- and TFB-TBOA-bound transporters in the OFS ^34^. When we lowered the temperature below 15 °C to slow <u>down</u> conformational transitions, we observed an additional S6 peak at ~ −218.5 ppm for RSMR-M385C-mFE (**Fig. S2**). Overall, the mFE resonances covered a range of 3.6 ppm. Ligands, mutations, and temperature affect the S1-S6 peak populations, suggesting that the resonances correspond to distinct structural states of the transporter and that their populations reflect the kinetic and equilibrium properties of the conformational ensemble. The three Glt_Ph_ protomers function independently ^30, 35^, and the labeling positions on the adjacent protomers are too distant to affect each other (**Fig. 1d**). Therefore, the ^19^F resonances report on the structural states of individual protomers.

**Figure 2.**
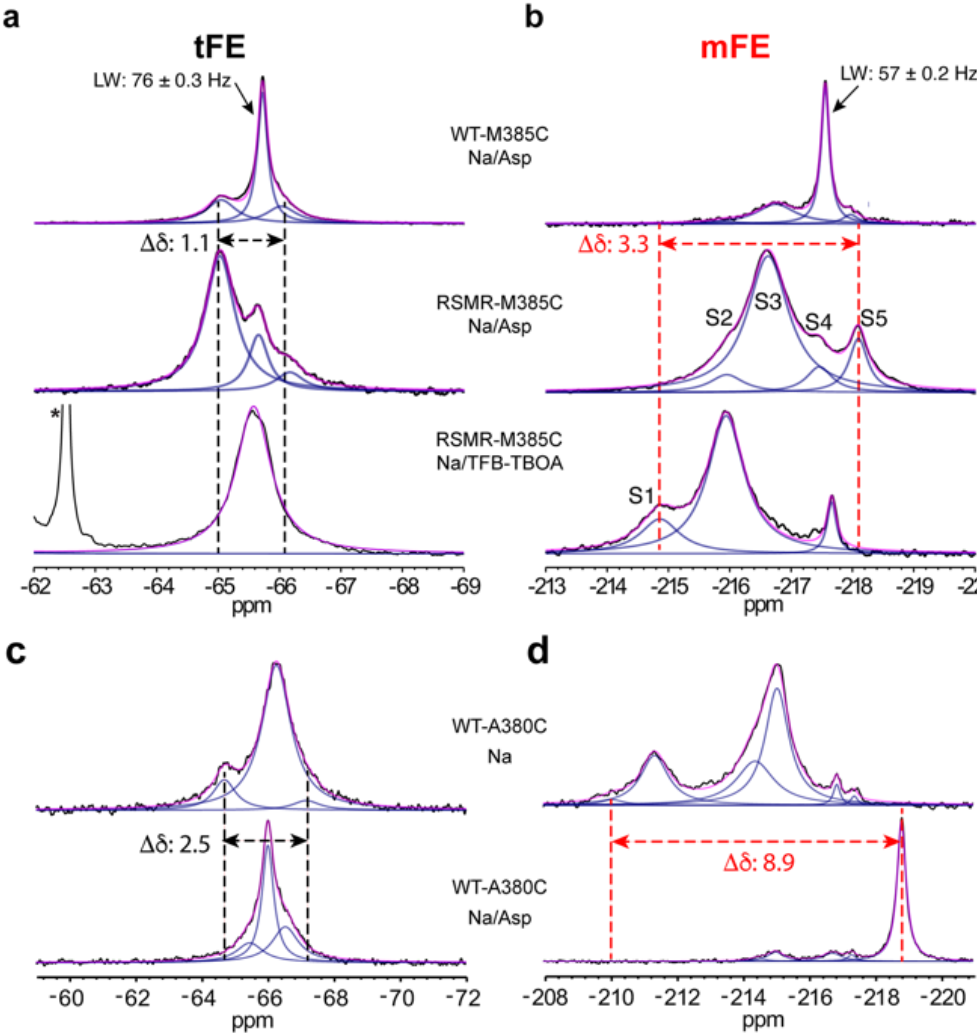
mFE-labeled Glt_Ph_ variants show wide chemical shift dispersion. ^19^F NMR spectra of Glt_Ph_-M385C (**a**, **b**) and Glt_Ph_-A380C (**c**, **d**) labeled with tFE (**a**, **c**) and mFE (**b**, **d**). From top to bottom: WT and RSMR mutant in 100 mM Na^+^ and 2 mM aspartate (Asp), the RSMR mutant in 100 mM Na^+^ and 0.6 mM TFB-TBOA (**a**, **b**), and WT in 400 mM Na^+^ and 100 mM Na^+^ and 2 mM aspartate (**c**, **d**). All spectra were recorded at 25 °C. Raw data are black, fitted spectra are pink, and deconvoluted Lorentzian peaks are blue. The asterisk denotes a signal from the CF_3_ group of TFB-TBOA. S1-S5 denote resolved resonances of Glt_Ph_-M385C-mFE. Δδ is the largest chemical shift difference observed for the labeling site. All ^19^F NMR spectra here and elsewhere were recorded at least twice on independently prepared protein samples, producing similar results.

We also compared tFE- and mFE-labeled WT-A380C and WT-A381C Glt_Ph_ mutants bound to Na^+^ ions only or Na^+^ ions and aspartate (**Fig. 2c, d** and **Fig. S4**). For both labeling sites, the mFE probe resulted in wider chemical shift dispersion. Strikingly, WT-A380C-mFE featured 7 peaks distributed over 8.9 ppm (**Fig. 2d**), demonstrating the tremendous potential of the probe. To our knowledge, this is the broadest chemical shift dispersion observed and the greatest number of protein states simultaneously resolved using ^19^F NMR.

### ^19^F-guided cryo-EM imaging to discover new conformational states

The ultrahigh environmental sensitivity of the mFE probe enables monitoring of how conformational equilibria are perturbed by the physicochemical environment, ligands, and mutations. Moreover, it can reveal hitherto uncharacterized structural states. For example, the ^19^F NMR spectra of WT-M385C-mFE and RSMR-M385C-mFE bound to the competitive blocker DL-*threo*-β-benzyloxyaspartic acid are dominated by distinct peaks, S4 for WT and S2 for RSMR (TBOA, **Fig. 3a**). Thus, unexpectedly, TBOA stabilizes the WT and RSMR transporters in different conformations. To establish whether these resonances correspond to OFS or IFS, we examined their solvent accessibility using paramagnetic relaxation enhancement (PRE). M385 is solvent-exposed in OFS but buried on the interface between the transport and scaffold domains in IFS (**Fig. 1e**) ^34, 36^. The addition of the soluble paramagnetic reagent Gd-DTPT-BMA significantly broadened the S4 but much less the S2 peak of the TBOA-bound RSMR mutant (**Fig. S5a**). Therefore, we assigned S4 to OFS and S2 to IFS. Similarly, for the Na^+^-bound WT, the paramagnetic reagent broadened S4 but not S2 or S3 (**Fig. S5b**), confirming the assignments of S4 and indicating that S3 is also IFS. The observation that the S2 and S3 peaks are intrinsically broader than S4 is also consistent with the faster transverse relaxation expected for a buried M385 site in these states.

**Figure 3.**
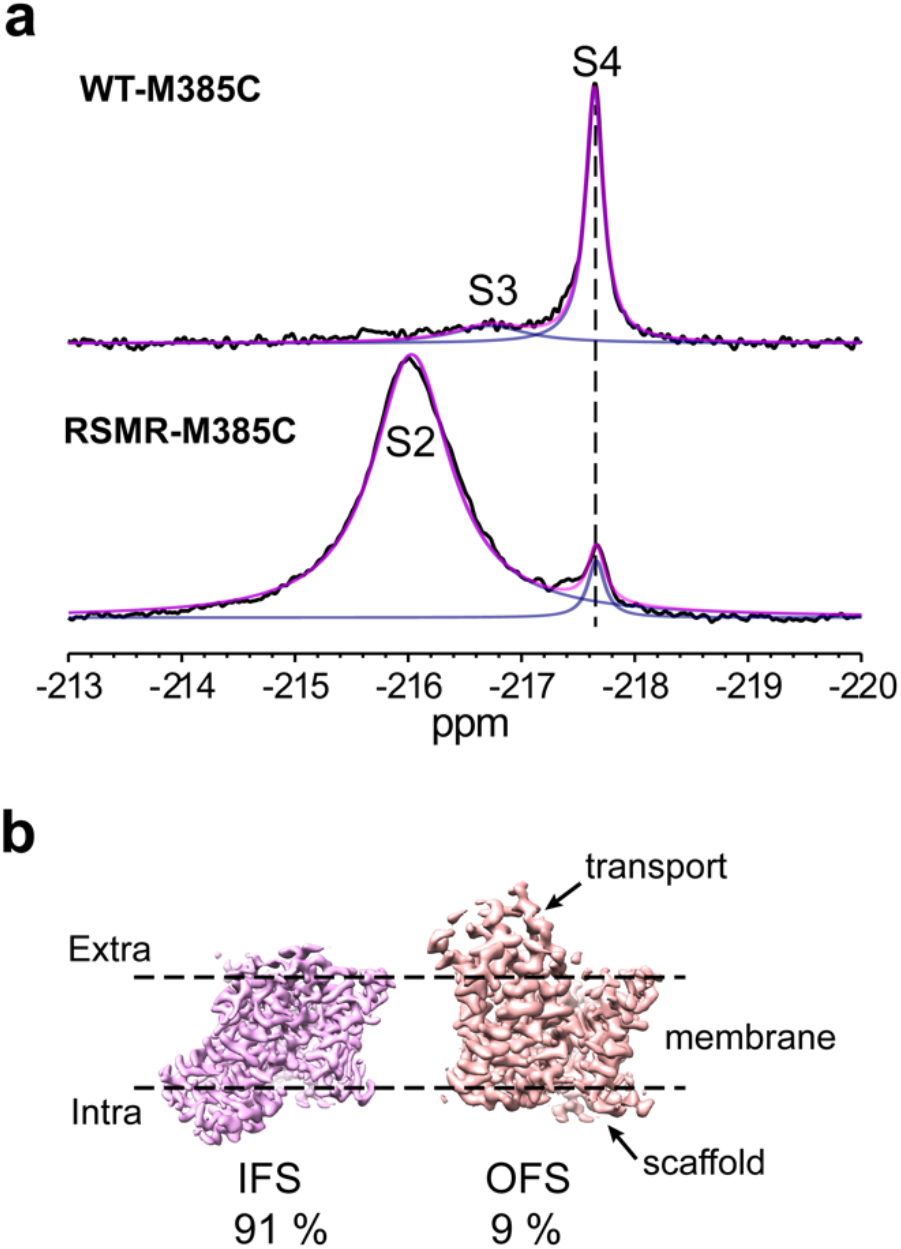
Identification and structural elucidation of new transporter conformations. (**a**) ^19^F NMR spectra of WT-M385C-mFE (upper panel) and RSMR-M385C-mFE (bottom panel) in the presence of 100 mM NaCl and 2 mM TBOA. The resonances occurring with similar chemical shifts to those in Fig. 2 are labeled S2, S3, and S4. (**b**) Cryo-EM density maps of TBOA-bound RSMR Glt_Ph_ mutant protomers in IFS (left) and OFS (right) structural classes. The corresponding populations are below the maps.

Our ^19^F NMR data revealed that the blocker TBOA stabilized the WT transporter in OFS, consistent with the published cryo-EM structure of this state ^34^. Unexpectedly, however, the blocker appeared to stabilize the RSMR mutant in an unknown IFS state. Motivated by this paradox, we imaged TBOA-bound RSMR by cryo-EM. Following particle alignment with imposed C3 symmetry, we performed symmetry expansion and 3D classification to sort multiple conformations of the transporter protomers ^34, 37^. We observed 91 % protomers in IFS with a wide-open substrate-binding site occupied by TBOA (**Fig. 3b**, **Fig. S6**, and **Table S1**; see *Methods* for data processing details). The remaining protomers were in OFS, similar to the TBOA-bound WT (RMSD of 0.646 Å). Because S2 and S4 are the only peaks in the RSMR-M385C-mFE NMR spectrum, we ascribe S2 to the highly populated open IFS and S4 to the minor OFS population observed by cryo-EM. Interestingly, the TBOA-bound WT populated a distinct S3 IFS (**Fig. 3a**). Consistently, a cryo-EM structure of the TBOA-bound WT transporter constrained in IFS by crosslinking showed a different, more closed conformation ^34^ than the structure we assigned to S2. Thus, the previously unobserved S2 state reveals a new modality of the ligand interaction with the transporter. While the physiological relevance of the S2 state is beyond the scope of this paper, it could represent an open-gate intermediate in the substrate release process.

### Synergistic use of ^19^F NMR and cryo-EM in exploring the conformation landscape

Under transport conditions in the presence of sodium and aspartate, ^19^F NMR spectra of RSMR-M385C-mFE showed that it populates a surprising number of conformational states, manifesting as resonances S1-S6 (**Fig. 2b, Fig. S2b** and **S2c**). Temperature modulated their populations so that, at 4 °C, we observed all states, with S6 being the most populated, while above 15 °C, S3 became dominant and S6 invisible (**Fig. S2**). Notably, the S6 peak decreased in amplitude and shifted to the left with increasing temperature, suggesting that it exchanges with another peak with rates approaching the NMR time scale of high μs to low ms. Solvent PRE at 15 °C, where S2 – S6 are well resolved, resulted in faster T_1_ relaxations of S4 and S5 but not S2, S3, and S6 resonances (**Fig. 4** and **Fig. S7**), suggesting that the mFE probe is exposed to the solvent for the former states and buried in the latter. The S1 peak is too small to evaluate by PRE. These results suggest that the RSMR mutant visits at least two OFS-like conformations, S4 and S5, and three different IFSs, S2, S3, and S6, exchanging slower than the NMR time scale. These assignments are consistent with the S2, S3, and S4 assignments described above for the TBOA-bound and Na^+^-bound transporters (**Fig. 3**, and **Fig. S6**)

**Figure 4.**
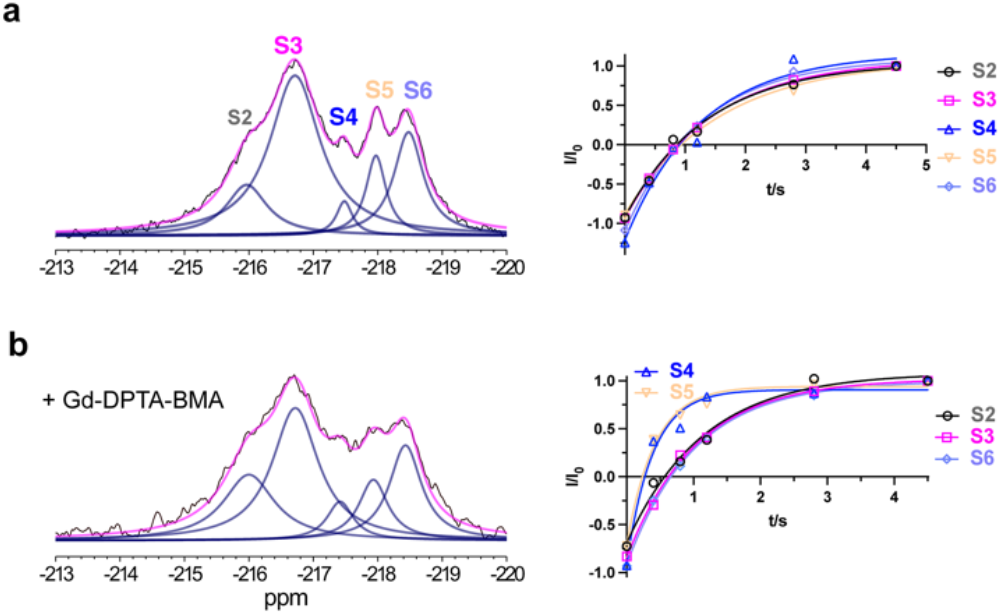
Assignment of IFS and OFS conformations based on solvent PRE effects. ^19^F NMR spectra (left) were recorded, and T_1_ relaxations (right) were measured in 300 mM NaCl and 2 mM Asp at 15 °C. Relaxation data were fitted to monoexponential functions (solid lines). The fitted T_1_ values are 1.36 ± 0.20, 1.41 ± 0.02, 1.26 ± 0.33, 1.56 ± 0.16, and 1.26 ± 0.09 s for S2, S3, S4, S5, and S6, respectively, in the absence of Gd-DPTA-BMA (**a**) and 1.15 ± 0.20, 1.04 ± 0.08, 0.39 ± 0.08, 0.42 ± 0.06, and 1.01 ± 0.06 s in the presence of 20 mM Gd-DPTA-BMA (**b**). The error bars, estimated as described in *Methods*, are too small to see.

The temperature-dependent population changes observed in ^19^F NMR experiments prompted us to ask whether cryo-EM can recapitulate them and inform the assignment of resonances to structural states. We flash-froze cryo-EM grids of RSMR-M385C-mET preincubated in 500 mM NaCl and 2 mM aspartate at 4, 15, and 30 °C. After data processing, 3D classification of protomers imaged at 4 °C revealed that ~87 % of them fell into IFS classes with variable orientations of the transport domain relative to the scaffold (**Fig. S8)**. The remaining ~13 % were in a previously described intermediate OFS (iOFS), in which the transport domain moves inward slightly compared to OFS. Based on similar transport domain positions, we grouped IFS classes into IFS-A and IFS-B ensembles with populations of 19 and 67 %, respectively (**Fig. 5 and Fig. S8a**). The main structural difference between them is that the transport domain packs more tightly against the scaffold in IFS-A but leans away and leaves a detergent-filled gap in IFS-B (**Fig. S8e**).

**Figure 5.**
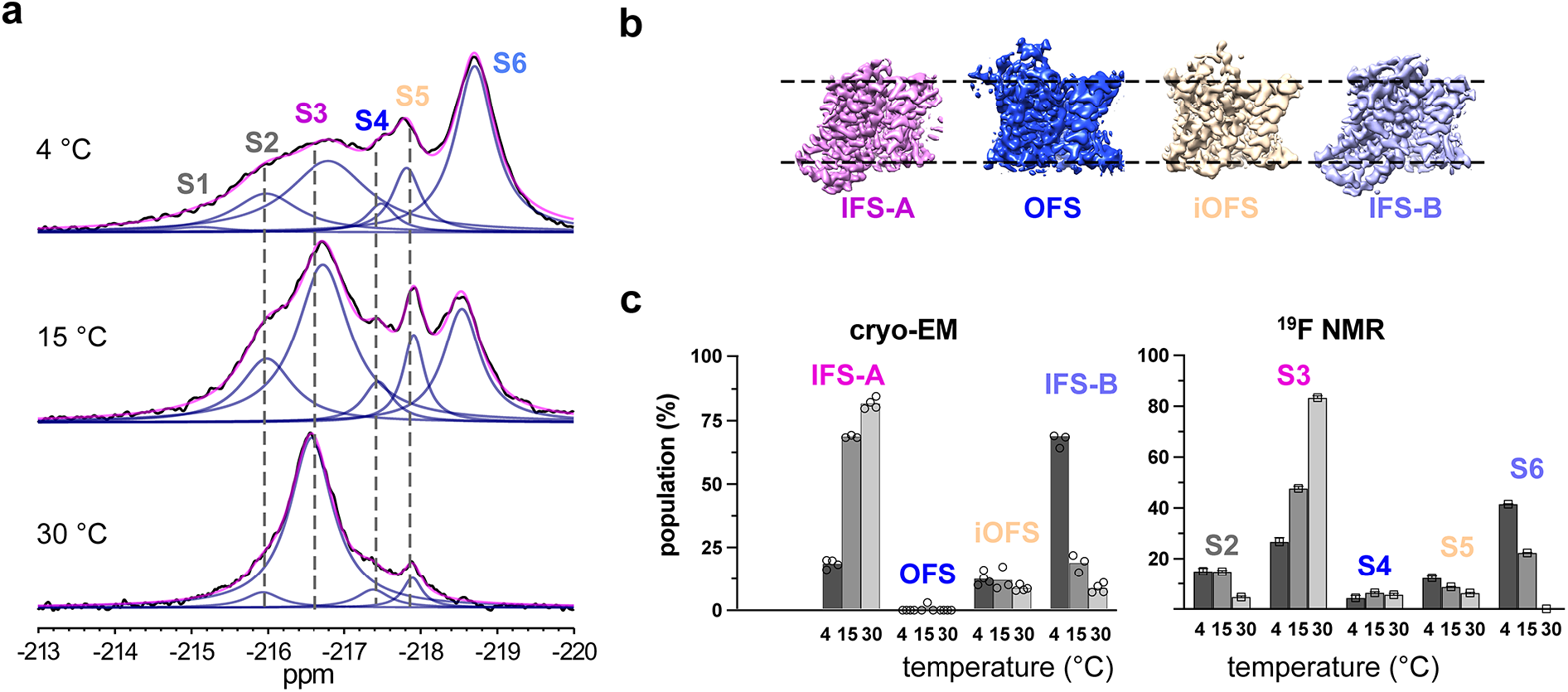
State populations observed in 3D classifications of protomers in cryo-EM parallel NMR measurements. (**a**) NMR spectra of RSMR-M385C-mFE were recorded in the presence of 500 mM NaCl and 2 mM Asp at 4 (top), 15 (middle), and 30 °C (bottom). (**b**) Representative maps of the four structural classes identified during cryo-EM particle 3D classifications of RSMR-M385C-mFE prepared at 4 °C. (**c**) The populations of protomers’ 3D classes in samples preincubated at 4, 15, and 30 °C (left). Open circles are results obtained using different T values and numbers of classes during 3D classifications in Relion. The state populations measured in ^19^F NMR experiments at the same temperatures are shown in the right panel. Errors are from multiple-peak deconvolutions of spectra in OriginPro.

Similar structures were observed crystallographically and termed “locked” and “unlocked”, respectively ^31^. The increasing temperature had little effect on the population of iOFS but led to a dramatic increase in IFS-A to ~ 82 % at 30 °C and a corresponding decrease in IFS-B populations to ~ 9 % (**Fig. 5c and Fig. S8b**).

The population shift from IFS-B to IFS-A at higher temperatures observed in cryo-EM parallels the increase of S3 and decrease of S6 in ^19^F NMR (**Fig. 5a**), strongly suggesting that S3 corresponds to IFS-A and S6 to IFS-B (**Fig. 5c**). Notably, the RSMS trimer crystallized at 4 °C with two protomers in the “unlocked” IFS-B and one protomer in the “locked” IFS-A ^31^, consistent with the higher IFS-B population observed by NMR and cryo-EM at 4 °C. S4, which corresponds to OFS, is lowly populated in RSMR but dominant in the WT transporter (**Fig. 2b** and **3a**). Consistently, cryo-EM didn’t identify the OFS in 4 °C data set. Instead, we only found ~ 2 % of particles in OFS in the 15 °C cryo-EM data set, reflecting the challenge in cryo-EM of detecting lowly populated states. S5 likely corresponds to iOFS because it is the only other observed structural state with the solvent-exposed 385C residue. While these assignments are plausible based on solvent exposure and the correlated population changes in the NMR and cryo-EM data, they remain speculative in the absence of direct structural data, such as we previously used to assign NMR signals to OFS from iOFS ^20^. The identities of the S1 and S2 resonances remain ambiguous. In particular, we did not observe a state in the cryo-EM classifications resembling the TBOA-bound RSMR structure, which populates the S2 peak (**Fig. 3**). Therefore, it is possible that one of the IFS conformations in the ensembles serendipitously shows a chemical shift similar to that of the TBOA-bound RSMR structure. The decrease in the S2 population at higher temperatures, thermodynamically similar to S6, suggests that it might correspond to one of the “unlocked” IFS-B subclasses.

## Discussion

Despite decades of efforts to develop biophysical methods to study protein dynamics, monitoring changes in membrane protein samples with more than two states remains challenging. The model protein in this study is a 134 kDa membrane transporter, Glt_Ph_, that undergoes complex conformational transitions to utilize transmembrane sodium ion gradients to drive the concentrative transport of aspartate. Resolving multi-conformational ensembles of such large membrane proteins using existing NMR methods is difficult. Our new mFE ^19^F probe takes advantage of the unique electronic properties of the monofluoroalkyl group leading to ultrahigh environmental sensitivity. It achieves chemical shift dispersion of ~9 ppm in the case of Glt_Ph_.

Relatively simple and inexpensive, ^19^F NMR using the new probe is an excellent tool for guiding high-resolution structural determinations by crystallography or cryo-EM. For example, ^19^F NMR showed that the transport blocker TBOA stabilizes the gain-of-function RSMR mutant of Glt_Ph_ in a different conformation than the WT. Consequent cryo-EM imaging led to the discovery of a novel conformation of the transporter in IFS featuring an open substrate gate and a new mode of inhibitor binding.

^19^F NMR resonances are difficult to assign to specific structural states because their chemical shifts are exquisitely sensitive to the local environment, indirectly reflecting global protein conformations. Previously, we showed that transition metal-mediated longitudinal PRE could assist with state assignments by evaluating the distance between the ^19^F label and a double histidine-coordinated metal ion. Here, we take advantage of enhanced line broadening and T_1_ relaxation due to the PRE effects of gadolinium compounds in solution to distinguish states with buried and exposed mFE probes. These measurements discriminate between OFS- and IFS-like states in WT Glt_Ph_ and its RSMR mutant.

Protein solutions are plunge-frozen for cryo-EM imaging in a process thought to preserve protein conformations, which can then be visualized by 3D classifications of the particles. Factors such as protein adsorption to the grids, surface tension and temperature changes during freezing may shift the conformation equilibrium. In addition, data processing, and parameters used in classifications can affect the obtained populations of 3D classes ^38^. Therefore, whether the class distributions reflect state populations in solutions is unclear. Here, for the first time, we evaluated the reliability of the state populations obtained from 3D classifications of EM-imaged particles. We found that the state populations determined by cryo-EM match well with those measured by ^19^F NMR. For example, the IFS and OFS populations of TBOA-bound RSMR were 95 and 5 % in ^19^F NMR and 91 and 9 % in cryo-EM, respectively. There was also good correspondence in complex ensembles of the aspartate-bound RSMR mutant preincubated at different temperatures. We observed a temperature-dependent correlated population increase of the main S3 resonance and the cryo-EM IFS-A structural ensemble and a corresponding decrease in the S6 resonance and the IFS-B ensemble.

Structural determination by cryo-EM and accurate state populations measured by ^19^F NMR provide a novel means to correlate the structural and thermodynamic properties of membrane proteins. For example, our results suggest that IFS-B has a dramatically lower enthalpy than IFS-A, resulting in the steep temperature dependence of the IFS-B to IFS-A transition. The lower IFS-B enthalpy must be associated with breaking the protein interface between the transport and scaffold domains and inserting detergent moieties into the gap. The detailed analysis of the phenomenon is beyond the scope of the paper. Nevertheless, our results show that even modest temperature changes can have profound effects on the observed conformational ensemble of a membrane protein, a feature characteristic, for example, of temperature-sensing ion channels^39^. ^19^F NMR is ideally suited to probe the temperature dependence of protein equilibria and might facilitate the study of the thermodynamics of membrane proteins-bilayer interactions.

^19^F NMR can detect conformations of RSMS-M385C-mEF present at only a few percent, such as the S4 peak, attributed to OFS. Such lowly populated states might be important functional intermediates, yet they are difficult to isolate during 3D classification of cryo-EM-imaged particles, especially when several conformations coexist. Thus, we only found OFS in the 15 °C cryo-EM dataset, where ^19^F NMR suggests its population is slightly higher than at 4 or 30 °C and where we collected the largest number of particles. Therefore, the ability of ^19^F NMR to visualize rare states provides a means to assess their populations and optimize conditions that can enrich them for high-resolution structural elucidation.

In summary, mFE chemical shift dispersion significantly exceeds that of tFE, enabling a more detailed and quantitative description of protein conformational ensembles and guiding cryo-EM imaging. Improved peak separation should also facilitate measurements of chemical exchange rates by methods such as EXSY ^20^ and saturation transfer, including CEST ^40^. Continuously expanding structural methodologies reveal that proteins populate diverse conformational ensembles. For example, a G protein-coupled adenosine A_2A_ receptor samples at least five distinguishable states during activation ^41^. Our model protein is similar in size to or larger than many physiologically important receptors, transporters, and ion channels. The new ^19^F probe should greatly facilitate mechanistic studies of such proteins.

## Materials and Methods

### Synthesis of S-(2-fluoroethyl-1,1,2,2-D_4_) *p*-toluenethiosulfonate

To a solution of potassium *p*-toluenethiosulfonate (678 mg, *TCI Chemical*) in dry acetonitrile (10 ml) was added 2-bromoethanol-1,1,2,2-d_4_ (240 μl, *Cambridge Isotope Laboratories*) under argon. The mixture was stirred at 60 °C overnight. The solvent was evaporated under reduced pressure, and the residue was dissolved in 50 ml ethyl acetate and washed with 0.5 M HCl aqueous solution (2 × 20 ml) and brine. The organic layer was dried over anhydrous Na_2_SO_4_ and then evaporated under reduced pressure to give S-(2-hydroxyethyl-1,1,2,2-D_4_) *p*-toluenethiosulfonate (600 mg, 85 % yield) as an oil. The product was directly used for the next step without further purification.

To a solution of S-(2-hydroxyethyl-1,1,2,2-D_4_) *p*-toluenethiosulfonate (480 mg, 2 mmol) in dry dichloromethane (5 ml) precooled to −70 °C, 1.5 equivalents of diethylaminosulfur trifluoride (3 ml of 1 M solution in dichloromethane, *Sigma-Aldrich*) were added dropwise under argon. After 10 min of stirring at −70 °C, the flask was warmed to room temperature for 30 min. The reaction was quenched with 0.5 ml of methanol. The reaction mixture was evaporated to dryness and then purified using a flash column (*RediSep Rf*) using 0-60 % petroleum ether/ethyl acetate. The major fraction was collected and evaporated to obtain the pure title compound (220 mg, 45 % yield). The identity of the compound was confirmed by NMR. ^1^H NMR (500 MHz, CDCl_3_) δ 7.82 (d, J = 8.4 Hz, 2H), δ 7.36 (d, J = 7.7 Hz, 2H), δ 2.46 (s, 3H); ^13^C NMR (125 MHz, CDCl_3_) δ 145.40, 141.85, 130.18, 127.24, 81.44-79.32 (J = 171.19 Hz), 34.88 (m), 31.84; ^19^F NMR (470 MHz, CDCl_3_) δ −215.25 (m).

### Synthesis of S-(2-fluoroethyl) *p*-toluenethiosulfonate

To a solution of potassium *p*-toluenethiosulfonate (620 mg, 2.7 mmol, *TCI Chemical*) in dry acetonitrile (10 ml) 1-bromo-2-fluoroethane (203 μl, *Accela ChemBio Inc*.) was added under argon. The mixture was stirred at 60 °C overnight. The solvent was removed under reduced pressure, and the residue was dissolved in 50 ml ethyl acetate and washed with 0.5 M HCl aqueous solution (2 × 20 ml) and brine. The organic layer was dried over anhydrous Na_2_SO_4_ and evaporated under reduced pressure. The residue was purified by flash column chromatography to give the pure title compound (503 mg, 86 % yield) as an oil. The identity of the compound was confirmed by NMR. ^1^H NMR (500 MHz, CDCl_3_) δ 7.82 (d, *J* = 8.2 Hz, 1H), 7.36 (d, *J* = 8.1 Hz, 1H), 4.61 (t, *J* = 6.3 Hz, 1H), 4.52 (t, *J* = 6.3 Hz, 1H), 3.30 (t, *J* = 6.3 Hz, 1H), 3.26 (t, *J* = 6.3 Hz, 1H), 2.46 (s, 3H); ^13^C NMR (126 MHz, CDCl_3_) δ 145.22, 141.72, 130.01, 127.11, 81.01 (d, *J* = 172.6 Hz), 35.38 (d, *J* = 22.7 Hz), 21.70; ^19^F NMR (471 MHz, CDCl_3_) δ −213.51 (tt, *J* = 46.9, 20.4 Hz).

### Protein expression, purification, and labeling

The single cysteine mutations A380C, A381C, and M385C were introduced into the Y215H/E219H cysteine-free Glt_Ph_ mutant (termed WT here for brevity), where the introduction of double histidine has been shown to not perturb the transport activity significantly ^20^. The M385C mutation was also introduced into the gain-of-function variant generated by additional R276S/M395R mutations (RSMR for brevity). All constructs were expressed, purified, and labeled as described, with modifications ^20, 36^. In brief, pBAD plasmids encoding constructs with C-terminal thrombin cleavage sites followed by (His)_8_-tag were transformed into *E. coli* DH10-B cells (*Invitrogen*). Cells were grown in LB media at 37 °C to an OD_600_ of 1.1. The temperature was then set to 30 °C and protein expression was induced by adding 0.2 % arabinose (*Goldbio*). Cells were grown for additional 16 hours. The cells were harvested by centrifugation and resuspended in 20 mM HEPES, pH 7.4, 200 mM NaCl, 1 mM L-asp, and 1 mM EDTA. The suspended cells were broken by an Emulsiflex C3 high-pressure homogenizer (*Avestin Inc*.) in the presence of 0.5 mg/ml lysozyme (*Goldbio*) and 1 mM phenylmethanesulfonyl fluoride (PMSF, *MP Biomedicals*). After centrifugation for 15 min at 5,000 g, the supernatant was centrifuged at 125,000 g for 50 min.

For mFE labeling, the membrane pellets were collected and solubilized in a buffer containing 20 mM HEPES, pH 7.4, 200 mM NaCl, 1 mM Asp, 10 mM 2-mercaptoethanol, 40 mM n-dodecyl-β-D-maltopyranoside (DDM, *Anatrace, Inc*.) for 2 hours at 4 °C. The mixture was centrifuged for 50 min at 125,000 g. The supernatant was diluted 3 times with buffer A (20 mM HEPES, pH 7.4, 200 mM NaCl, 1 mM Asp) and incubated with Ni-NTA resin (*Qiagen*) for 1 hour at 4 °C. The resin was washed with 6 volumes of Buffer A supplemented with 1 mM DDM and 25 mM imidazole and proteins were eluted in buffer A supplemented with 1 mM DDM and 300 mM imidazole. EDTA was added to the collected protein fractions to a final concentration of 0.5 mM. The protein was concentrated to ~10 mg/ml using concentrators with 100 kDa MW cutoff (*Amicon*). Protein concentration was determined by UV absorbance at 280 nm using the extinction coefficient of 57,400 M^−1^ cm^−1^ and MW of 44.7 kDa (protomer). Two molar equivalents of TsSCD_2_CD_2_F or TsSCH_2_CH_2_F, prepared as stock solutions in DMSO, were added to the protein samples followed by incubation at 4 °C for 2 hours or room temperature for 1 hour. Thrombin was added and incubated overnight at room temperature to cleave the (His)_8_-tag.

For tFE labeling, the membrane pellet was solubilized in a buffer containing 20 mM HEPES, pH 7.4, 200 mM NaCl, 1 mM Asp, 40 mM DDM, and 2 mM 2,2’-dithiodipyridine (DTDP, *Sigma Aldrich*). After binding to Ni-NTA resin, the protein-resin slurry was first washed with 5 volumes of Buffer A supplemented with 1 mM DDM. Trifluoroethanethiol (2 mM, *Sigma Aldrich*) was added, and the slurry was incubated with mixing at 4 °C overnight. The resin was washed with buffer A supplemented with 1 mM DDM and 25 mM imidazole and then eluted with buffer A supplemented with 1 mM DDM and 300 mM imidazole. The (His)_8_-tag was cleaved using thrombin at room temperature overnight.

Both mFE- and tFE-labeled proteins were further purified by size exclusion chromatography (SEC) using a Superdex 200 Increase 10/300 GL column (*GE Healthcare Life Sciences*) in a buffer containing 20 mM HEPES, pH 7.4, 50 mM KCl, and 1 mM DDM. NaCl (100 mM) was added to the protein fractions immediately. The protein was concentrated and supplemented with additional NaCl and ligands as needed. The labeling efficiency was quantified using 1D NMR and found to be quantitative (**Fig. S1a**)

### Transport activity assay

Unlabeled and mFE-labeled WT Glt_Ph_ were concurrently reconstituted into liposomes, and initial rates of ^3^H Asp uptake were measured as previously described ^42^. Briefly, liposomes were prepared from a 3:1 (w/w) mixture of *E. coli* polar lipids and egg yolk phosphatidylcholine (*Avanti Polar Lipids*) in 20 mM HEPES/Tris, pH 7.4, containing 200 mM KCl and 100 mM choline chloride. Liposomes were destabilized by the addition of Triton X-100 at detergent to lipid ratio of 0.5:1 (w/w). Glt_Ph_ proteins were added at a final protein-to-lipid ratio of 1:2,000 (w/w) and incubated for 30 min at room temperature. Detergents were removed with Bio-Beads™ SM-2 resin (*Bio-Rad*) via two incubations at room temperature, one overnight incubation at 4 °C, and two more incubations at room temperature. The proteoliposomes were concentrated to 50 mg/mL and flash-frozen in liquid N_2_. On the day of the experiment, liposomes were thawed and extruded through 400 nm filters. The uptake reaction was started by diluting the proteoliposomes 100-fold into reaction buffer containing 20 mM HEPES/Tris pH 7.4, 200 mM KCl, 100 mM NaCl, 1 μM ^3^H-L-Asp (*PerkinElmer*), and 0.5 μM valinomycin. Uptake was measured at 2 minutes at 35 °C. Reactions were quenched by adding 10 volumes of cold buffer containing 20 mM HEPES/Tris pH 7.4, 200 mM KCl, and 100 mM LiCl. Uptakes of the mutants were normalized to the WT protein in each experiment. Each data point is an average of at least two technical replicates, and the data are composed of the results from three independent liposome reconstitutions.

### ^19^F NMR spectroscopy

^19^F NMR spectra were collected on Bruker Advance III HD 500 MHz spectrometer equipped with a TCI ^1^H-^19^F/^13^C/^15^N triple resonance cryogenic probe (*Bruker Instruments*) tuned for ^19^F. For mFE-labeled proteins, 50 μM 2-fluoroethanol and 10 % D_2_O were added to the sample and used as a chemical shift reference (−224.22 ppm) and a lock signal, respectively. Typically, 160 μl of the transporter solution at a final concentration of 100 - 250 μM (protomer) was loaded into a 4 mm Shigemi tube (Shigemi Co., Ltd). 1D ^19^F NMR spectra were recorded using the standard ZG pulse in the Bruker pulse library, with 2,096 points recorded and a spectral width (SW) of 40 ppm. The carrier frequency was set at - 220 ppm. The recycle delay was set to 1.5 and 0.9 s for deuterated and protonated mFE, respectively, except when otherwise indicated. The scan numbers were set according to protein and salt concentrations and temperatures between 2 and 30 K to achieve a satisfactory signal-to-noise ratio. For tFE-labeled proteins, 50 μM trifluoroacetic acid was added to the samples and used as a chemical shift reference (−76.55 ppm), the carrier frequency was set at −70 ppm, and the recycle delay was set to 0.6 s. All the spectra were recorded without decoupling.

All 1D ^19^F NMR spectra were processed using MestReNova 12.0.0 software (*Mestrelab Research*). The free induction decay signals were zero-filled to 8,192 points and Fourier-transformed after applying a 20 Hz exponential window function. The spectra were baseline-corrected, and the peaks were manually picked. The initial values of peak heights and linewidths were manually set such that their sum approximated the original spectrum. Simulated annealing was used for iterative fitting. During fitting, linewidths were constrained between 20 and 800 Hz, peak positions were constrained within 5% variation, and the peak shapes were set to Lorentzian. The reported linewidth values were obtained by subtracting the line-broadening value of 20 Hz from the fitted linewidth. To test the reliability of the fitting procedure and the reproducibility of the resulting parameters, we recorded spectra of three independently prepared RSMR-M385C-mFE samples in 300 mM NaCl and 2 mM Asp. The obtained fitted values of chemical shifts, linewidths, and state populations showed small standard deviations (**Fig. S3a-c, e**). Additionally, we exported MestReNova-processed spectra and fitted them using the Multiple Peak Fit tool in OriginPro 2019 software (*OriginLab*), which also reported small fitting errors (**Fig. S3d**).

^19^F longitudinal relaxation times (*T*_1_) were measured by the inversion recovery method. The recycle delays were set to 5 and 2.5 s for deuterated and protonated mFE probes, respectively. The spectra were processed in MestReNova, imported into OriginPro (*OriginLab*), and globally fitted using the Global Fit Tool patch. The peaks were picked, and initial estimates of their chemical shifts, linewidths, and heights were set manually. The chemical shift and linewidth values for each peak were held fixed between all spectra in the relaxation series but varied relative to other peaks during fitting. The peak intensities and areas were normalized using values obtained after the longest delay. *T*_1_ relaxation times were estimated by fitting the relaxation plots to single exponential functions *I* = *I*_0_(1 − 2*exp*(−*t*/*T*_1_)), where *I* is the normalized peak intensity, and *t* is the relaxation time.

To explore whether probe deuteration sharpens the resonance peaks as expected, we compared the spectra of deuterated and protonated mFE attached to WT-M385C in the presence of saturating Na^+^ ions, where we observed well-resolved S3 and S4 peaks (**Fig. S9**). Consistent with deuterium reducing the ^19^F-^1^H dipole relaxation and peak splitting due to ^19^F-^1^H spin-spin coupling, deuterated mFE produced sharper peaks than protonated mFE with a more pronounced effect on the intrinsically sharper S4 peak (**Fig. S9**). However, the longer T_1_ relaxation times (**Fig. S9**), increasing the recycle delay and reducing sensitivity, offset the benefits of deuteration. If hardware enabling ^1^H decoupling during ^19^F detection is available the protonated mFE probe might be preferable.

### Cryo-EM data collection

To prepare cryo-grids, 3.5 μL of labeled Glt_Ph_ protein (4.5 mg/mL) was applied to a glow-discharged QuantiFoil R1.2/1.3 300-mesh gold grid (*Quantifoil Micro Tools GmbH*) and incubated for 2 min under 100 % humidity at the desired temperatures. Following incubation, grids were blotted for 3 s at 0 blot force and plunge-frozen in liquid ethane using a Vitrobot Mark IV (*Thermo Fisher Scientific*). Cryo-EM imaging data were acquired on a Titan Krios microscope (*Thermo Fisher Scientific*) operated at 300 kV with a K3 Summit direct electron detector (*Gatan, Inc*.). Automated data collection was carried out in superresolution mode with a magnification of 105,000 x, which corresponds to a calibrated pixel size of 0.852 Å on the specimen and 0.426 Å for superresolution images. An energy slit width of 20 eV was used throughout the collection. For the TBOA-bound RSMR sample, movies were collected using Leginon ^43^ at a total dose of 52.88 e^−^/Å^2^ distributed over 48 frames (1.102 e^−^/ Å^2^/frame) with an exposure time of 2.40 s (50 ms/frame) and a defocus range of 1.3 − 2.0 μm. A total of 8,100 movies were collected. For the Asp-bound RSMR grids made at 4 °C, movies had a total dose of 50.94 e^−^/Å^2^ distributed over 48 frames (1.061 e^−^/ Å^2^/frame) with an exposure time of 2.40 s (50 ms/frame) and a defocus range of 1.3 – 1.5 μm. A total of 5267 movies were collected. For the Asp-bound RSMR grids prepared at 15 °C, movies had a total dose of 58.57 e^−^/Å^2^ distributed over 50 frames (1.171 e^−^/ Å^2^/frame) with an exposure time of 2 s (40 ms/frame) and a defocus range of 0.9 – 1.9 μm. A total of 14132 movies were collected. For the Asp-bound RSMR grids prepared at 30 °C, movies had a total dose of 56.04 e^−^/Å^2^ distributed over 40 frames (1.401 e^−^/ Å^2^/frame) with an exposure time of 1.6 s (40 ms/frame) and a defocus range of 1.3 – 1.6 μm. A total of 5823 movies were collected.

### Image processing

The frame stacks were motion corrected using MotionCorr2 ^44^ with 2x binning, and contrast transfer function (CTF) estimation was performed using CTFFIND4.1 ^45^. Further processing steps were carried out using RELION 3.0.8 or 3.1.0 and cryoSPARC 3.0 or 3.2 ^46–47^. Particles were picked from micrographs using the Laplacian-of-Gaussian (LoG) picker, aiming for ~2000 picks per micrograph. These particles were extracted using a box size of 300 pixels with 4x binning and imported into cryoSPARC. Following one round of 2D classification to remove artifacts, the particles underwent 3 rounds of heterogeneous refinement in C1 using eight classes. Seven classes were noise volumes created by one iteration of *ab initio*, and one was an unmasked 3D volume obtained from a complete *ab initio* run. The particles were converted to Relion format using PyEM ^48^ and re-extracted at full box size. These particles were reimported into cryoSPARC and underwent one round of nonuniform (NU) refinement using C3 symmetry ^49^. These particles were converted back to Relion format for Bayesian polishing with parameters obtained using 5,000 random particles ^50^. The polished particles were reimported into cryoSPARC and subjected to one round of NU refinement in C3 with both local and global CTF refinement options turned on. The particles were again polished in Relion using an expanded box size of 384 pixels, reimported into cryoSPARC, and subjected to one round of NU refinement with C3 symmetry and local and global CTF refinement options turned on. The particles were then C3 symmetry-expanded and subjected to a focused 3D classification in Relion. The local mask was generated by UCSF Chimera^51^ using a combination of OFS (chain A of PDB model 2NWX) and IFS (chain A of PDB model 3KBC). The T value was set between 10 and 40 depending on the job. The 3D class populations were calculated by dividing the particle number in the individual class or a group of similar classes by the total number of C3 symmetry-expanded particles. The particles from the individual classes or combined similar classes were imported separately into cryoSPARC and subjected to local refinement using the mask and map obtained from the most recent NU refinement.

### Model building and refinement

Crystal or cryo-EM structures were used as initial models and docked into the density maps using UCSF Chimera. The models were first real-space refined in PHENIX^52^. Misaligned regions were manually rebuilt, and missing side chains and residues were added in COOT^53^. Models were iteratively refined with applied secondary structure restraints and validated using MolProbity^54^. To cross-validate models, all atoms in the refined models were randomly displaced by an average of 0.3 Å, and each resulting model was refined against the first half-map obtained from processing. The FSC between the refined models and the first half-maps was calculated and compared to the FSC of the other half-maps. The resulting FSC curves were similar, showing no evidence of overfitting (**Fig. S10**). The structural figures were prepared in UCSF Chimera and PyMOL (DeLano Scientific).

## Acknowledgments

The authors thank Dr. M. Goger and Dr. S. Bhattacharya for their help with setting up NMR. We thank Dr. B. Wang, Dr. A. Paquette, and Dr. W. Rice at the NYU cryo-EM facility for help with electron microscopy data collection. We also thank W. Eng and E. Wu for assistance with protein expression, and Dr. Z. Shen for assistance with chemical synthesis. This work was supported by NIH grants R37NS085318 and R01NS064357 (O.B.), R37AG019391 and R35 GM136686 (D.E.), and S10OD016320 (C.B.). O.B., D.E., >and C.B. are members of the New York Structural Biology Center (NYSBC), which is supported in part by NIH Grant P41 GM118302 (CoMD/NMR: Center on Macromolecular Dynamics by NMR Spectroscopy), ORIP/NIH facility improvement grant CO6RR015495 and NIH grant S10OD018509.

## Author contributions

Y.H., D.E., and O.B. designed the experiments; Y.H. performed the NMR experiments; Y.H. and O.B analyzed the data; C.B. assisted with NMR experimental data collection and analysis; Y. H and W. Z conducted the chemical synthesis; Y. H., K. R. and B. Q. performed cryo-EM imaging, analyzed data, and refined molecular models; K. R. performed transport activity assays; Y.H., K. R., O.B. and D.E. wrote the manuscript.

## Table of Contents (TOC)

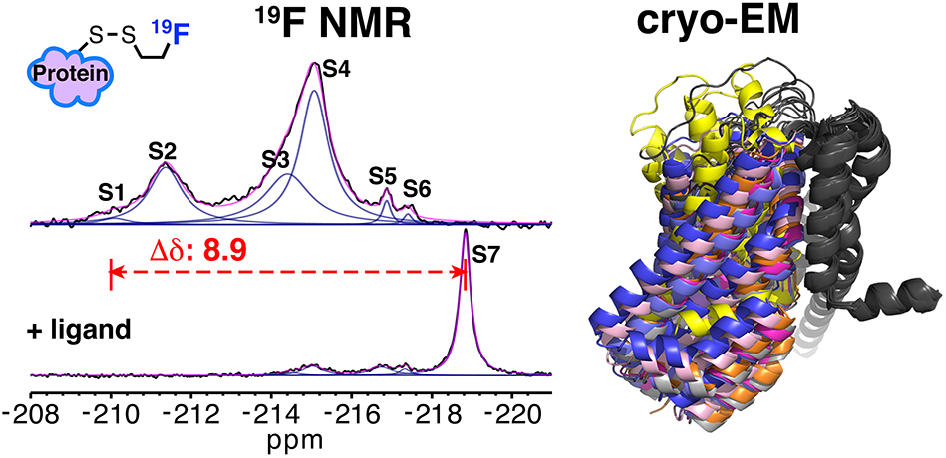

