## Supplemental information for "An environmentally ultrasensitive fluorine probe to resolve protein conformational ensembles by ^19^F NMR and cryo-EM"

### Contents:

#### Supplementary figures

**Fig. S1**. Labeling efficiency and $^3$H-aspartate uptake assay. (P2)

**Fig. S2**. Temperature-dependent spectra of tFE- and mFE-labeled RSMR-M385C. (P3)

**Fig. S3**. Reproducibility of NMR peak deconvolution (P4)

**Fig. S4**. Chemical shift dispersion of tFE- and mFE-labeled WT-A381C (P5)

**Fig. S5**. $^{19}$F peak assignment based on solvent PRE effects. (P6)

**Fig. S6**. Cryo-EM structures of TBOA-bound RSMR mutant. (P7-P8)

**Fig. S7**. Global peak deconvolution of paramagnetic $T_1$ relaxation (P9)

**Fig. S8**. Temperature-dependent Na/Asp-bound RSMR structural ensembles. (P10-P11)

**Fig. S9**. Relaxation properties of deuterated and protonated mFE probes. (P12)

**Fig. S10**. FSC of Map and model validations. (P13)

**Table S1**. Cryo-EM data collection, reconstruction, and model refinement statistics. (P14-P15)

**NMR characterization of the labeling compounds.** (P16-P18)

**Extended Data**

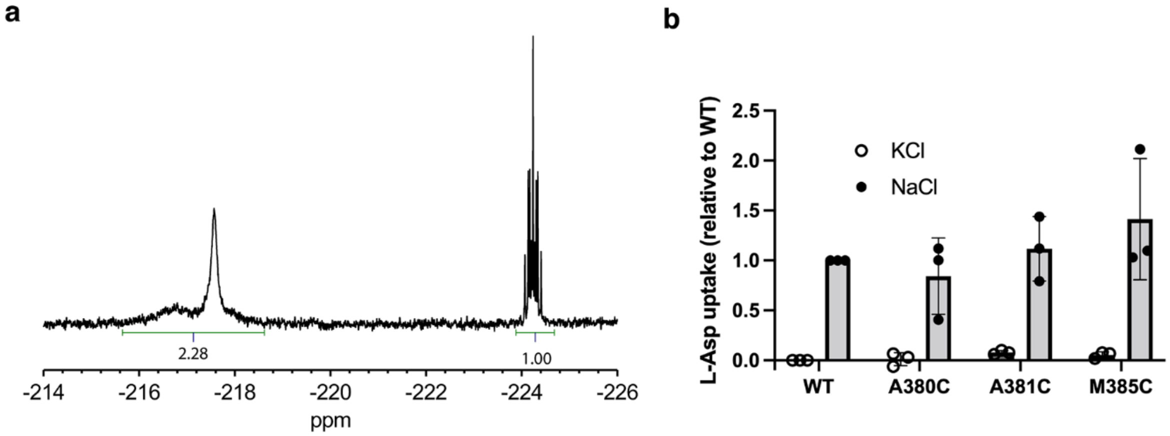


**Figure S1. Quantification of the labeling efficiency and activity of labeled proteins.** **(a)** Spectra of 220 µM WT-M385C-mFE $Glt_{Ph}$ and 100 µM 2-fluoroethanol standard were recorded in 20 mM HEPES, pH 7.4, 1 mM DDM, 100 mM NaCl, 2 mM Asp, 10 % $D_2O$. The recycle delay was set to 10 s to ensure accurate quantification. The 2-fluoroethanol and mFE signals were integrated (green lines and corresponding areas below the peaks), and the amount of the attached mFE label was determined as 228 µM. **(b)** $^3H$-Asp uptake assay of WT $Glt_{Ph}$ and mFE-labeled single-cysteine mutants (see Methods). The initial $^3H$-Asp uptake rates were measured at 2 min for mFE-labeled $Glt_{Ph}$ variants relative to an unlabeled cysteineless WT $Glt_{Ph}$. Data were normalized to WT uptake in the presence and absence of sodium gradients (closed circles and open circles, respectively). Experiments were repeated in triplicates on independently prepared protein samples.

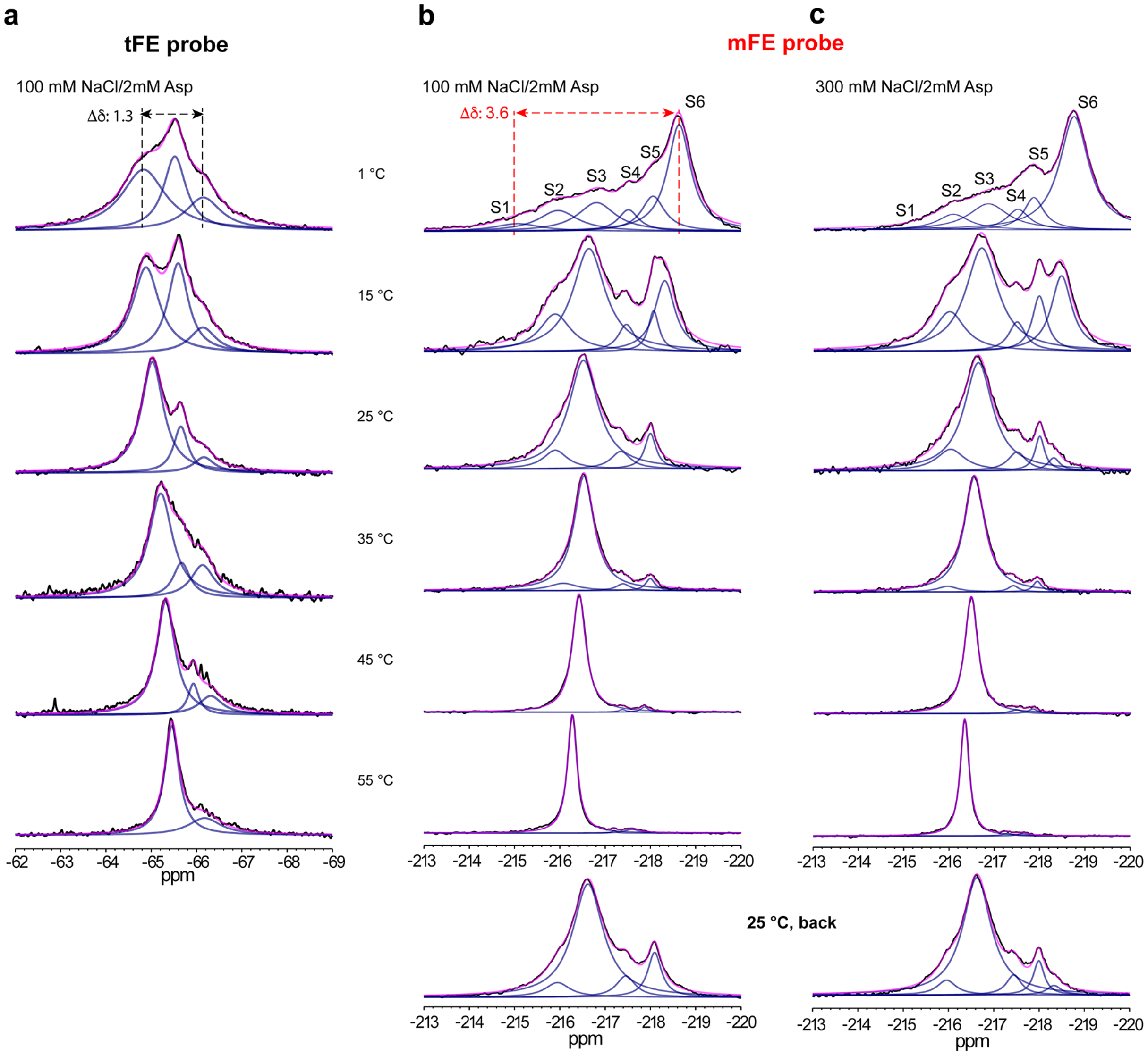


**Figure S2. Temperature dependence of $^{19}$F NMR spectra of tFE- (a) and mFE-labeled (b, c) RSMR-385C.** Spectra were recorded in the presence of 100 mM NaCl and 2 mM Asp (**a, b**) and 300 mM NaCl and 2 mM Asp (**c**) at indicated temperatures. Spectra recorded after cooling the sample from 55 back to 25 °C are shown at the bottom of panels b and c. $\Delta\delta$ indicates the largest observed chemical shift difference in ppm.

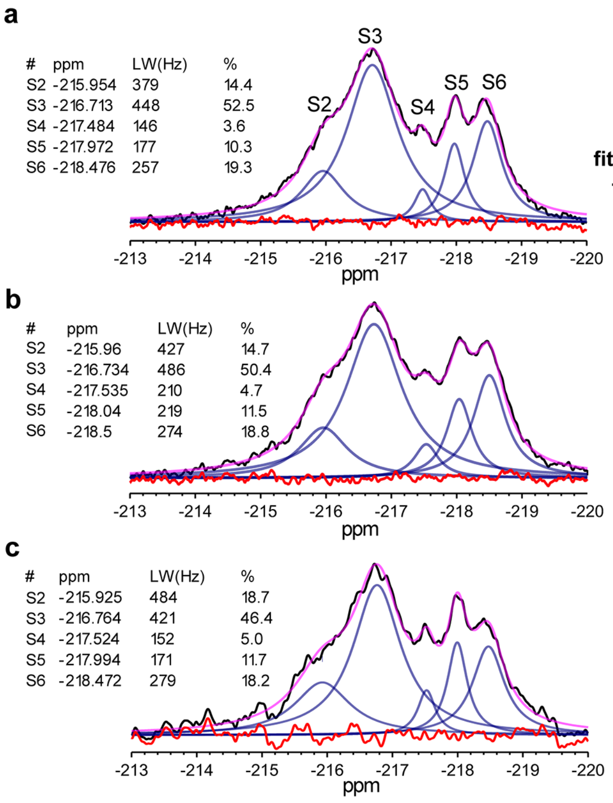

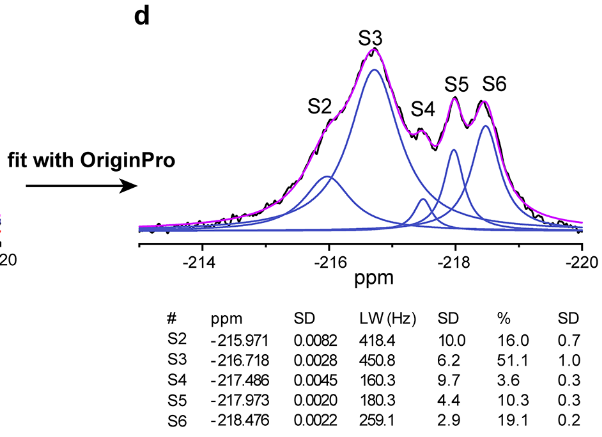


**e**

| Peak | Chemical shift (ppm) | Linewidth (Hz) | Population (%) |
|---|---|---|---|
| S2 | -215.95 ± 0.02 | 420 ± 53 | 16 ± 2 |
| S3 | -216.74 ± 0.03 | 452 ± 33 | 48 ± 3 |
| S4 | -217.51 ± 0.03 | 169 ± 35 | 4.9 ± 0.2 |
| S5 | -218.00 ± 0.03 | 189 ± 26 | 11.6 ± 0.1 |
| S6 | -218.48 ± 0.02 | 270 ± 12 | 18.5 ± 0.4 |

**Figure S3. Reproducibility of NMR peak deconvolution. (a-c),** Peak deconvolutions of $^{19}F$ NMR spectra of three biological repeats using MestReNova. The protein samples at 15 °C were in a buffer containing 20 mM HEPES, pH 7.4, 1 mM DDM, 300 mM NaCl, and 2 mM Asp. The original raw spectra, fitted spectra, deconvoluted Lorentzian peaks, and fitting residuals are in black, pink, light blue, and red, respectively. The chemical shifts (ppm), linewidths (LW, Hz), and peak populations (%) are shown to the left of the spectra. (**d),** Peak deconvolution of the spectrum in (**a)** using the Multiple Peak Fit tool in OriginPro. The fitting results with standard deviations (SD) are below the spectrum. (**e**) Mean fitted values from (**a-c**) with standard deviations.

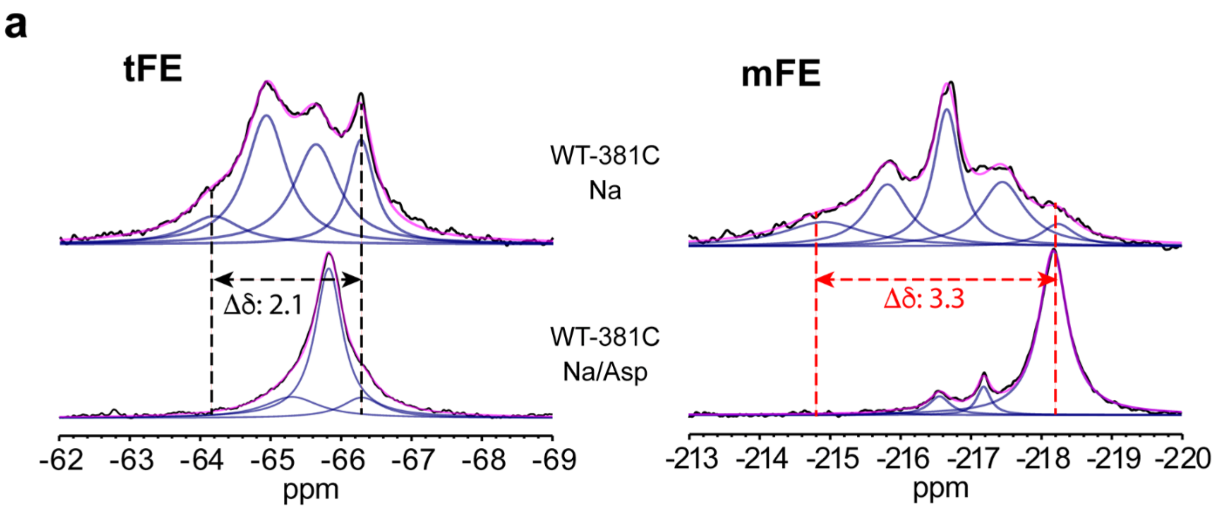


**Figure S4. Comparison of chemical shift dispersion of tFE- and mFE-labeled WT-A381C**. $^{19}F$ NMR spectra of WT-A381C labeled with tFE (left) and mFE (right). Shown are spectra in the presence of 400 mM $Na^{+}$ (upper panel) and 100 mM $Na^{+,}$ and 2 mM Asp (lower panel). All spectra were recorded at 25 °C. $\Delta\delta$ indicates the largest observed chemical shift difference in ppm.

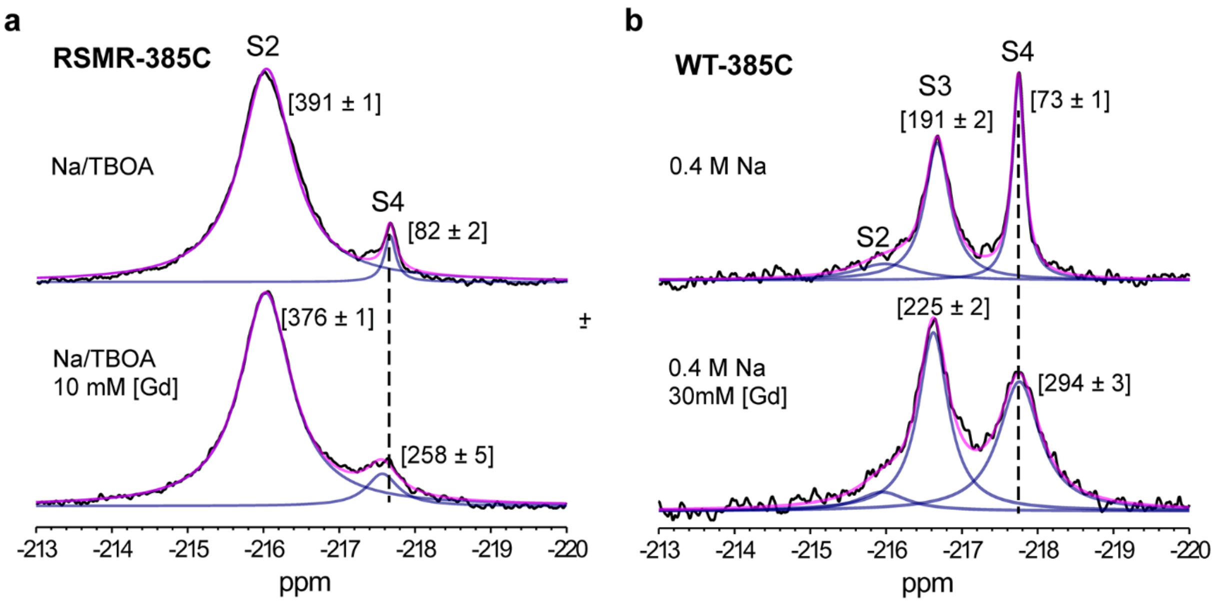


**Figure S5. $^{19}$F peak assignment based on solvent PRE effects.** (**a**) $^{19}$F NMR spectra of RSMR-M385C-mFE in the presence of 100 mM NaCl and 2 mM TBOA without (top) and with (bottom) 10 mM Gd-DPTA-BMA. (**b**) $^{19}$F NMR spectra of WT-M385C-mFE in the presence of 400 mM NaCl without (top) and with (bottom) 30 mM Gd-DPTA-BMA. All spectra were recorded at 25 °C. The numbers next to the peaks correspond to the linewidths and errors from peak deconvolution.

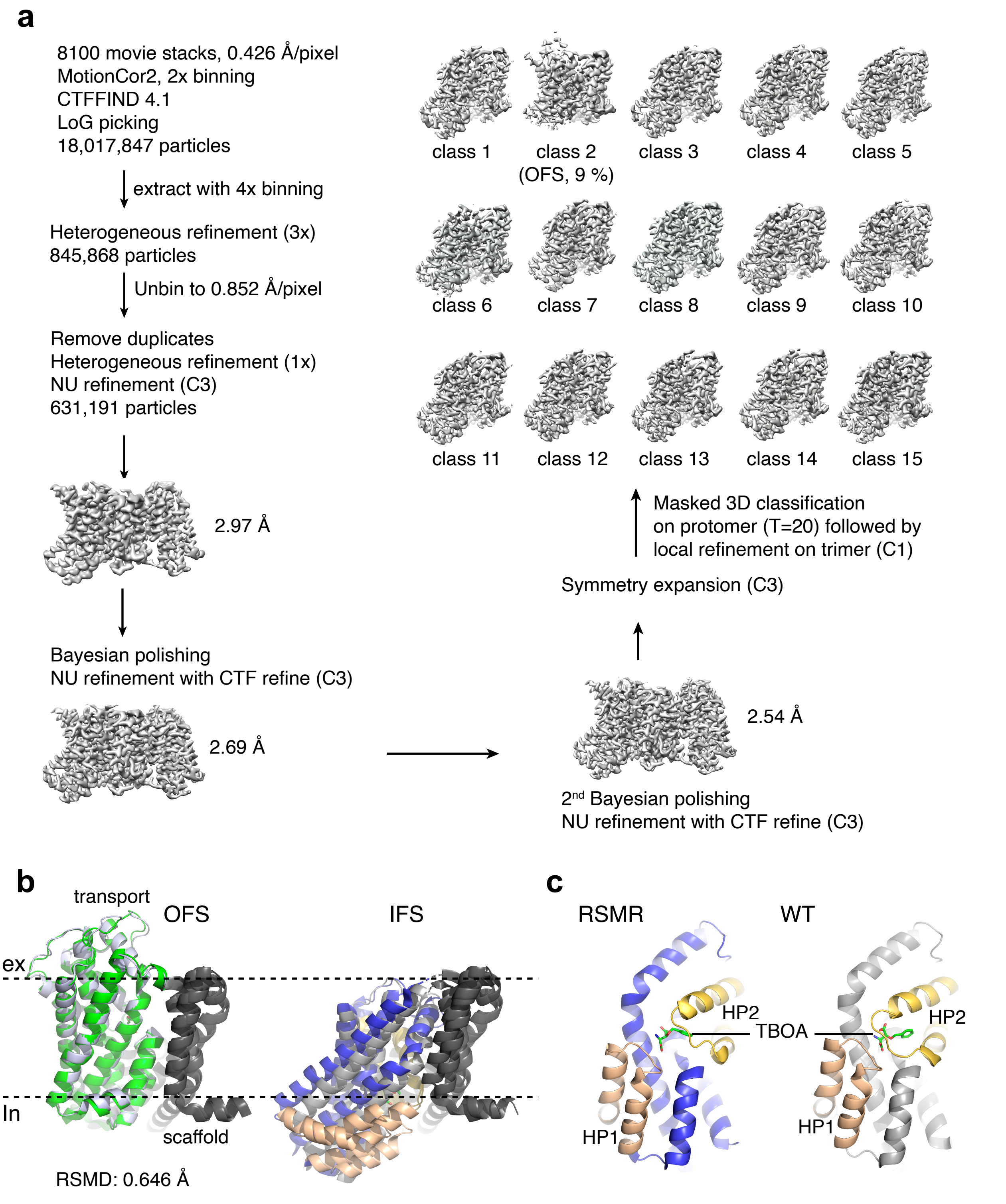


**Figure S6. Cryo-EM structures of TBOA-bound RSMR mutant**. (**a**) Data processing scheme. Only Class 2 is OFS (9 %), and all other classes are IFSs. (**b**) RSMR mutant in OFS (left) and IFS (right). The TBOA-bound RSMR mutant (green) and WT Glt$_{Ph}$ (silver,

PDB code 6X17) in OFS were superimposed over the entire protomers with RMSD shown below the structure. The TBOA-bound RSMR mutant refined from all IFS classes (blue) and WT $Glt_{Ph}$ cross-linked in IFS (silver, PDB code 6X16) were aligned on the scaffold trimerization residues 140-215. The scaffold domain is dark gray. (**c**) Transport domains of the RSMR (left) and cross-linked WT (right) transporters in IFS. TBOA is shown as sticks. The helical hairpins (HP) 1 and 2 are colored wheat and gold, respectively.

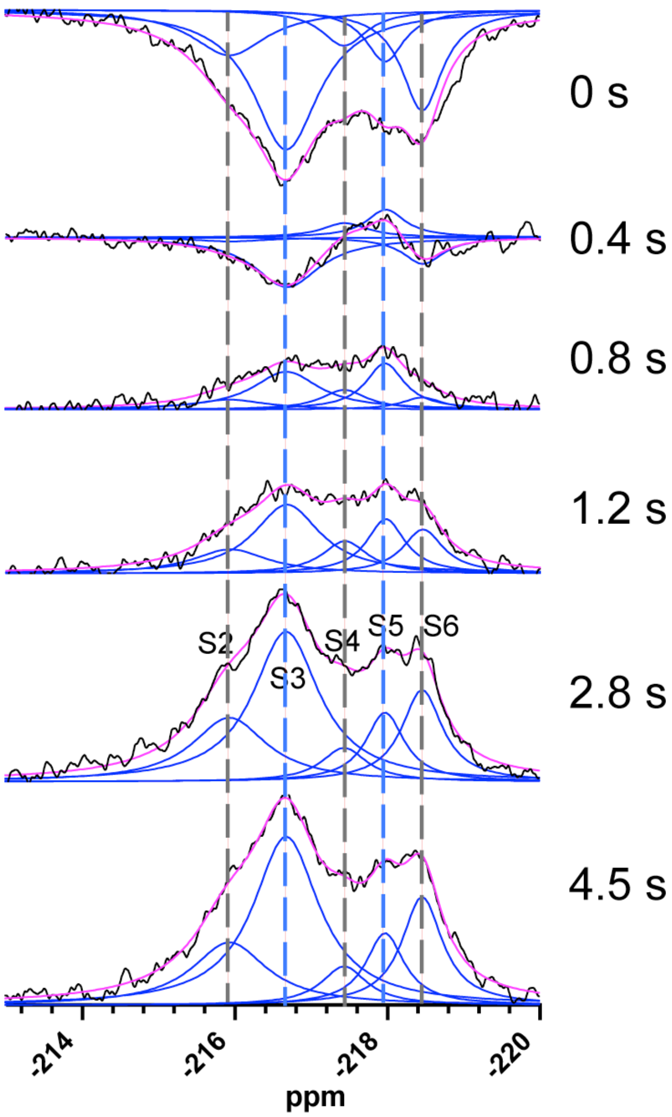


**Figure S7. Peak deconvolution of the paramagnetic $T_1$ relaxation of the Asp-bound RSMR mutant**. The spectra were acquired after the relaxation delays shown on the right. The spectra deconvolution was carried out globally using OriginPro with chemical shifts and linewidths as global fitted parameters for the entire dataset and peak heights as local parameters for each spectrum.

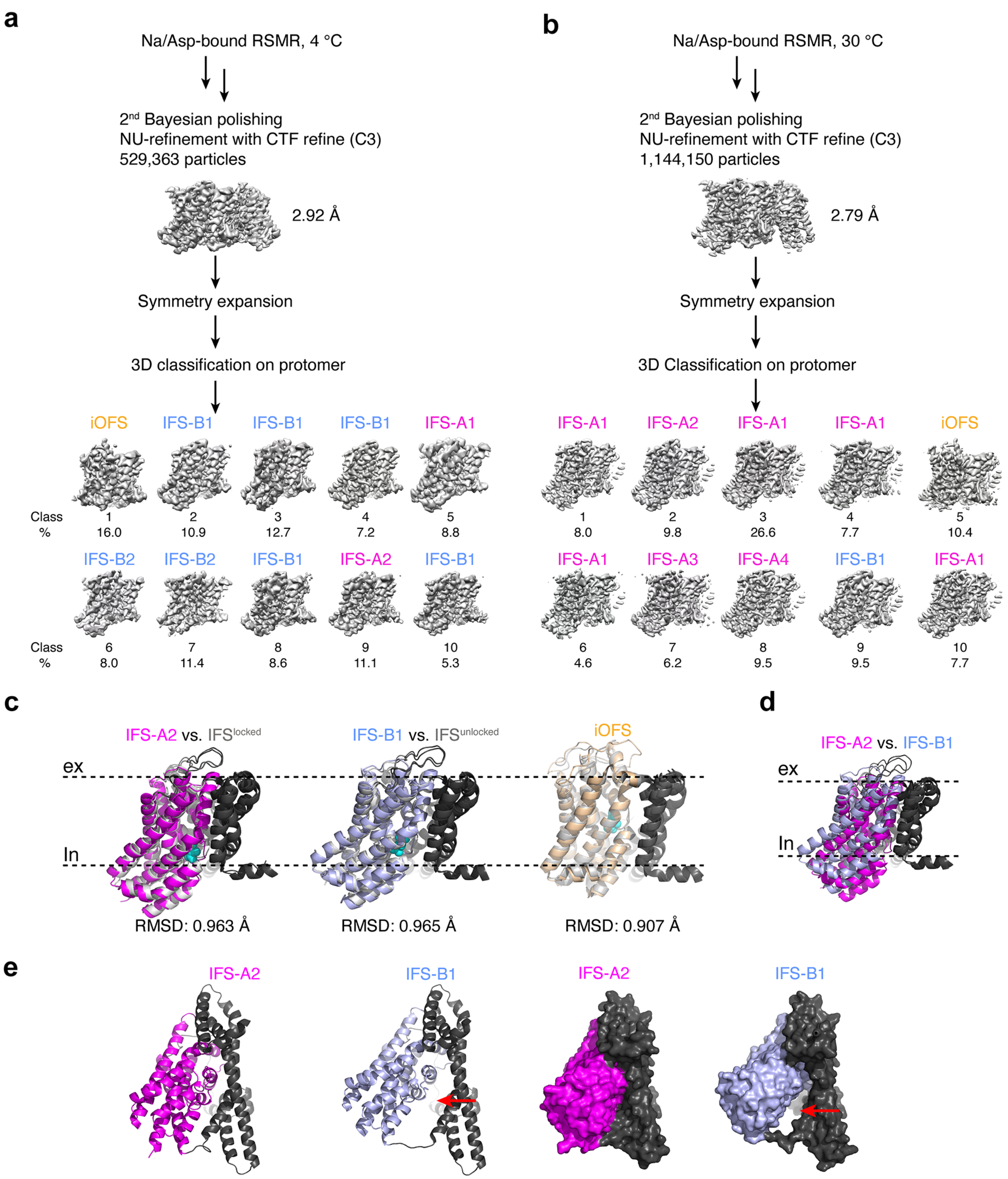


**Figure S8. Temperature-dependent structural ensembles of the aspartate-bound RSMR mutant**. Cryo-EM data processing schemes for grids prepared at 4 (**a**) and 30 °C (**b**). Examples of 3D classification results are shown with class populations below the density maps. IFS structural classes IFS-A and IFS-B comprise several subclasses, IFS-A1-4 and IFS-B1-2, showing small movements of the transport domains. (**c**)

Superpositions of an aspartate-bound IFS-A2 (magenta, left), IFS-B1 (light blue, middle), and iOFS (wheat, right) with the previously reported structures (silver): “locked” IFS (PDB code 3KBC), “unlocked” IFS (4X2S), and iOFS (6UWL), respectively. The scaffold domain is colored dark gray. Corresponding RMSDs are below the structures. (**d**) A superposition of IFS-A2 (magenta) and IFS-B1 (blue), viewed in the membrane plane. (**e**) Cytoplasmic views of IFS-A2 and IFS-B1 in cartoon (left) and surface (right) representations. The red arrows indicate the detergent-filled gap between the transport and scaffold domains. Structures were aligned on the scaffold domain.

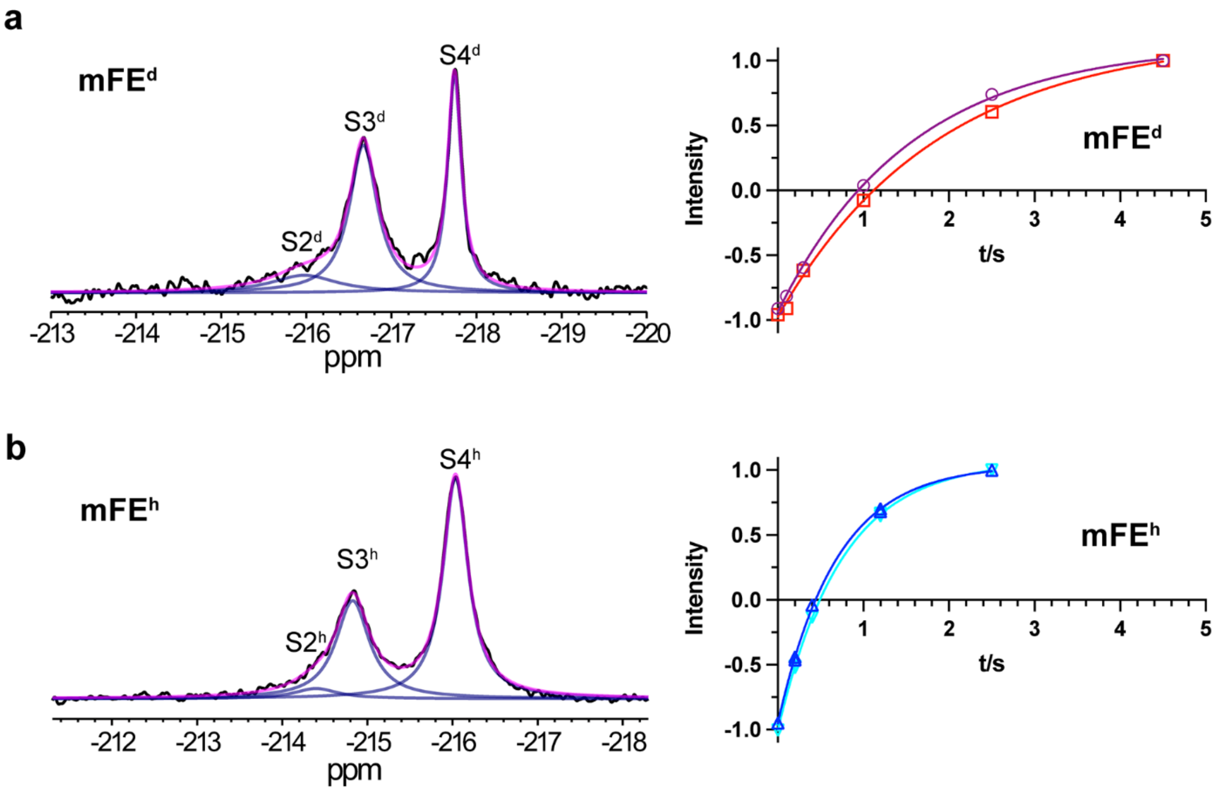


**Figure S9**. **Relaxation properties of deuterated (mFE$^{d}$) and protonated (mFE$^{h}$) mFE probes.** (**a**) $^{19}$F NMR spectra of mFE$^{d}$- labeled WT-385C in the presence of 400 mM NaCl (left) and $T_1$ relaxation plots (right) of the S3$^{d}$ (purple circles) and S4$^{d}$ (red squares). Peak deconvolution gave linewidth of S3$^{d}$ and S4$^{d}$ as 196 $\pm$ 3.3 Hz and 81 $\pm$ 0.6 Hz, respectively. Solid lines of the plots are least-square fits to mono-exponential equations, with time constants of 1.55 $\pm$ 0.08 s and 1.87 $\pm$ 0.16 s for the S3$^{d}$ and S4$^{d}$ peak, respectively. (**b**) $^{19}$F NMR spectra of mFE$^{h}$- labeled WT-385C in the presence of 400 mM NaCl (left) and $T_1$ relaxation plots (right) of the S3$^{h}$ (blue triangles) and S4$^{h}$ (cyan invert triangles). Peak deconvolution gave linewidth of S3$^{h}$ and S4$^{h}$ as 236 $\pm$ 2.1 Hz and 150 $\pm$ 0.6 Hz, respectively. Solid lines of the plots are least-square fits to mono-exponential equations, with time constants of 0.68 $\pm$ 0.02 s and 0.74 $\pm$ 0.01 s for the S3$^{d}$ and S4$^{d}$ peak, respectively. All measurements were conducted at 25 °C. The error bars on the intensities obtained from the global peak deconvolution are too small to see.

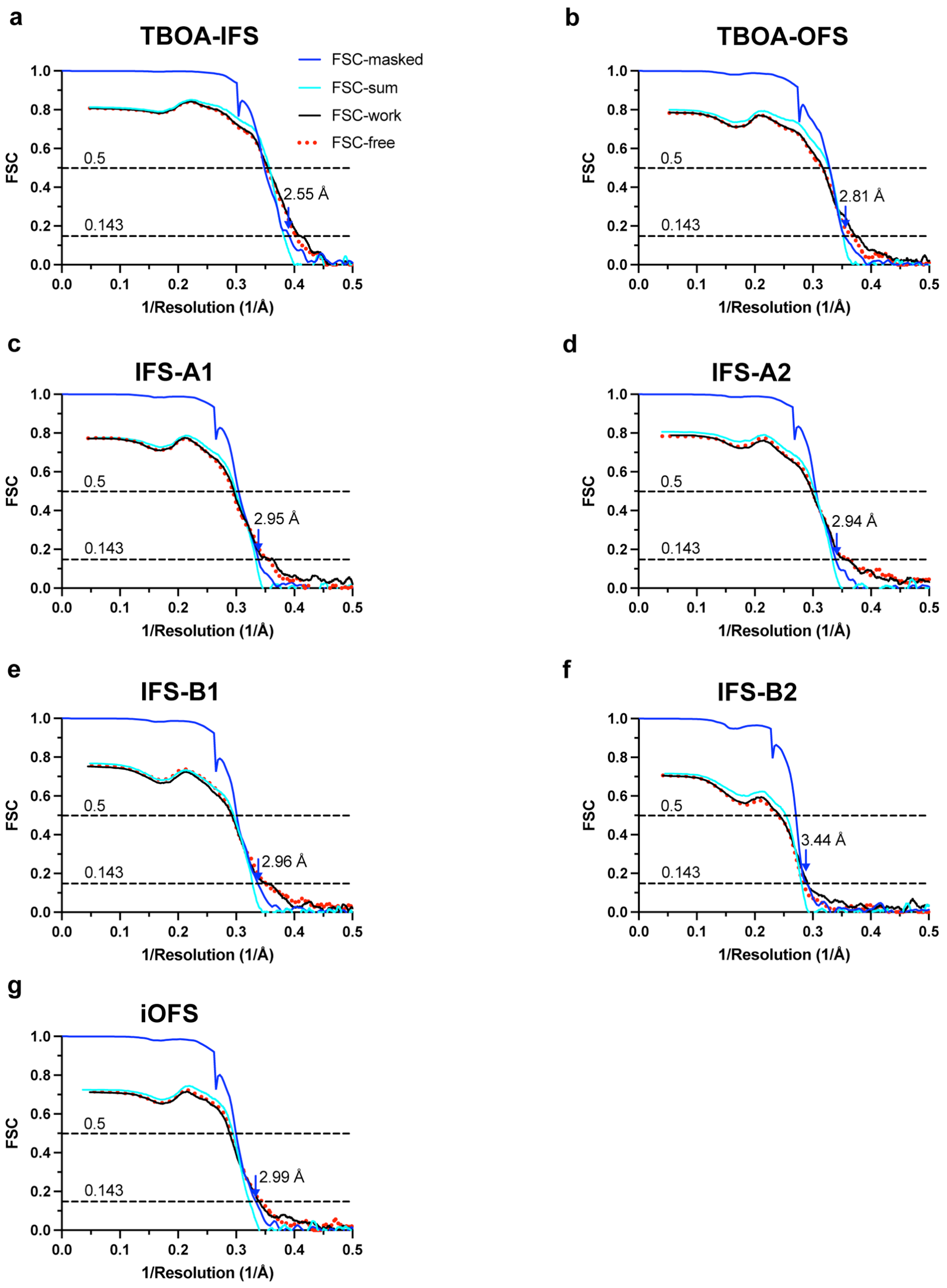


**Figure S10**. **Map and model validations.** (**a**) Fourier Shell Correlation (FSC) curves for the density maps and map-model validations in the following states: TBOA-bound IFS (**a**), TBOA-bound OFS (**b**), Asp-bound IFS-A1 (**c**), Asp-bound IFS-A2 (**d**), Asp-bound IFS-B1 (**e**), Asp-bound IFS-B2 (**f**) and Asp-bound IFS-iOFS (**g**). Shown are the FSC curves for the density maps (blue) and the FSC curves for the refined models versus full maps (cyan) and half maps for cross-validation (black lines and red dots). Dashed lines correspond to the FSC values of 0.5 (up) and 0.143 (bottom), respectively.

**Extended Data Table S1. cryo-EM data collection, reconstruction, and model refinement statistics**

| | RSMR-TBOA (25 °C) [a] | | RSMR-Asp (30 °C) [a,b] | | | | RSMR-Asp (4 °C) [a,c] |
|---|---|---|---|---|---|---|---|
| **Data collection and processing** | | | | | | | |
| Magnification | 105,000 | | 105,000 | | | | 105,000 |
| Voltage (kV) | 300 | | 300 | | | | 300 |
| Electron exposure (e–/Å$^2$) | 52.88 | | 50.94 | | | | 56.04 |
| Defocus range (μm) | 1.3-2.0 | | 1.3-1.5 | | | | 1.3-1.6 |
| Pixel size (Å) | 0.4260 | | 0.4260 | | | | 0.4260 |
| Micrographs (no) | 8,100 | | 5267 | | | | 5823 |
| Initial particles (no.) | 18,017,847 | | 12,174,463 | | | | 20,578,986 |
| Final particles (no.) | 631,191 | | 396,595 | | | | 1,144,150 |
| **Reconstruction** | | | | | | | |
| | IFS (EMD-26482) (PDB 7UG0) | OFS (EMD-26489) (PDB 7UGJ) | IFS-A1 (EMD-26487) (PDB 7UGD) | IFS-A2 (EMD-26497) (PDB 7UGV) | IFS-B1 (EMD-26498) (PDB 7UGX) | iOFS (EMD-26504) (PDB 7 UH3) | IFS-B2 (EMD-26505) (PDB 7UH6) |
| Particle no. | 1,731,338 | 162,235 | 272,276 | 335,559 | 326,008 | 356,418 | 147,516 |
| Symmetry imposed | C1 | C1 | C1 | C1 | C1 | C1 | C1 |
| Map resolution (Å) | 2.55 | 2.81 | 2.95 | 2.94 | 2.96 | 2.99 | 3.38 |
| FSC threshold | 0.143 | 0.143 | 0.143 | 0.143 | 0.143 | 0.143 | 0.143 |
| Map resolution range (Å) | 1.859 – 31.177 | 1.842 – 36.514 | 1.972 – 40.064 | 1.829 – 40.061 | 1.858 – 40.105 | 1.969 – 39.643 | 2.152 – 47.367 |
| **Refinement** | | | | | | | |
| Initial model used (PDB code) | 6X16 | 6X17 | 3KBC | 3KBC | 4X2S, chain B | 6UWL | 4X2S, chain B |
| Model resolution (Å) | 2.8 | 3.1 | 3.3 | 3.3 | 3.4 | 3.4 | 3.9 |
| FSC threshold | 0.5 | 0.5 | 0.5 | 0.5 | 0.5 | 0.5 | 0.5 |
| Map sharpening *B* Factor (Å$^2$) | -59.1 | -68.5 | -93.0 | -94.8 | -92.7 | -98.3 | -88.8 |
| Model composition | | | | | | | |
| Non-hydrogen atoms | 3,126 | 3,095 | 3,079 | 3,107 | 3,108 | 3,096 | 3,095 |
| Protein residues | 418 | 416 | 415 | 418 | 416 | 416 | 416 |
| Ligands | 3 | 3 | 3 | 3 | 3 | 3 | 3 |
| *B* factors (Å$^2$) | | | | | | | |
| Protein | 88.05 | 106.67 | 122.57 | 123.11 | 114.55 | 126.59 | 130.34 |
| Ligand | 96.22 | 114.65 | 125.45 | 121.84 | 135.18 | 141.52 | 145.43 |
| R.m.s. deviations | | | | | | | |
| Bond lengths (Å) | 0.002 | 0.002 | 0.002 | 0.003 | 0.002 | 0.003 | 0.003 |
| Bond angles (°) | 0.376 | 0.420 | 0.460 | 0.574 | 0.469 | 0.59 | 0.612 |
| Validation | | | | | | | |
| MolProbity score | 1.01 | 1.17 | 1.00 | 1.12 | 1.15 | 1.19 | 1.34 |
| Clashscore | 2.34 | 3.79 | 2.21 | 3.29 | 3.61 | 4.09 | 6.13 |
| Poor rotamers (%) | 0 | 0 | 0 | 0 | 0 | 0 | 0 |
| Ramachandran plot | | | | | | | |
| Favored (%) | 99.03 | 99.27 | 99.27 | 99.27 | 98.54 | 98.05 | 98.29 |
| Allowed (%) | 0.97 | 0.73 | 0.73 | 0.73 | 1.46 | 1.95 | 1.71 |
| Disallowed (%) | 0 | 0 | 0 | 0 | 0 | 0 | 0 |

[a] Chamber temperature for making grids. [b] Additional maps corresponding to IFS-A3 and IFS-A4 were deposited in https://www.wwpdb.org as EMD-29514 and EMD-29522, respectively, without models. [c] IFS-A1, IFS-A2, IFS-B1, IFS-B2, and iOFS conformations

were identified and the corresponding maps were deposited as EMD-29544, EMD-29545, EMD-29543, EMD-26505, and EMD-29542, respectively, but a structural model was built only for IFS-B2 for the 4 °C data.

### NMR characterization of the labeling compounds

**Figure S11.** $^1$H NMR spectrum of $TsSCD_2CD_2F$

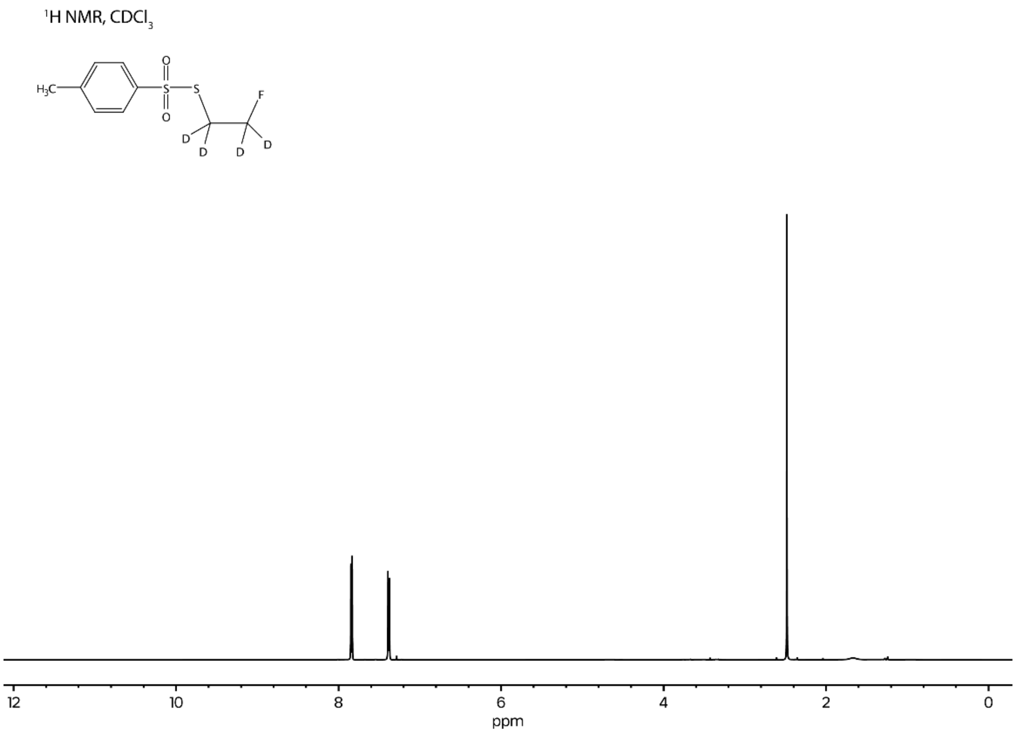


**Figure S12.** $^{13}$C NMR spectrum of $TsSCD_2CD_2F$

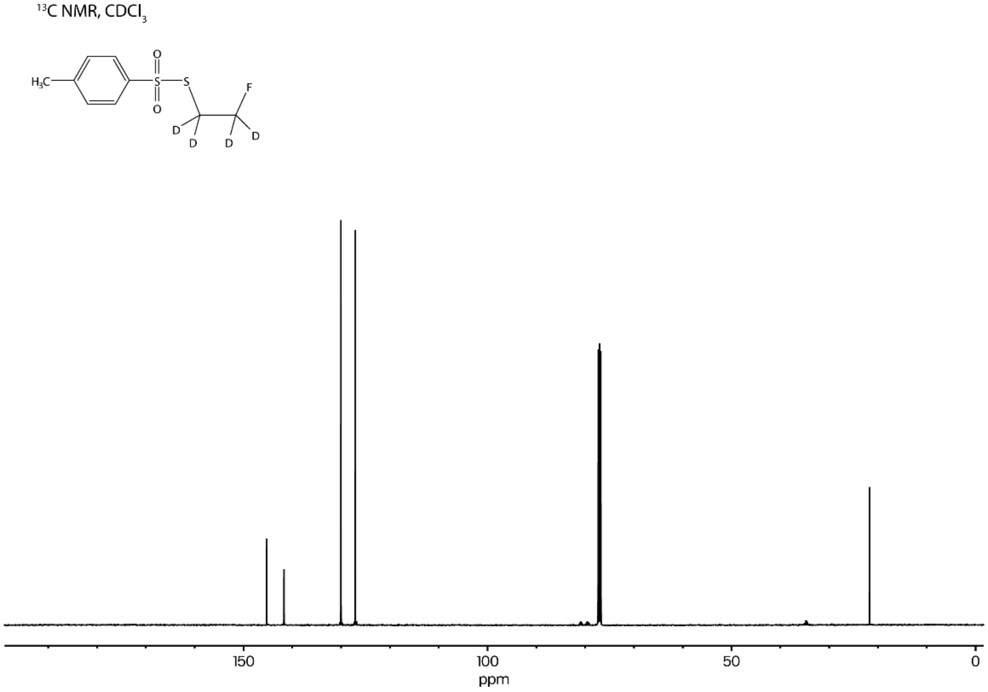

**Figure S13.** $^{19}F$ NMR spectrum of $TsSCD_2CD_2F$

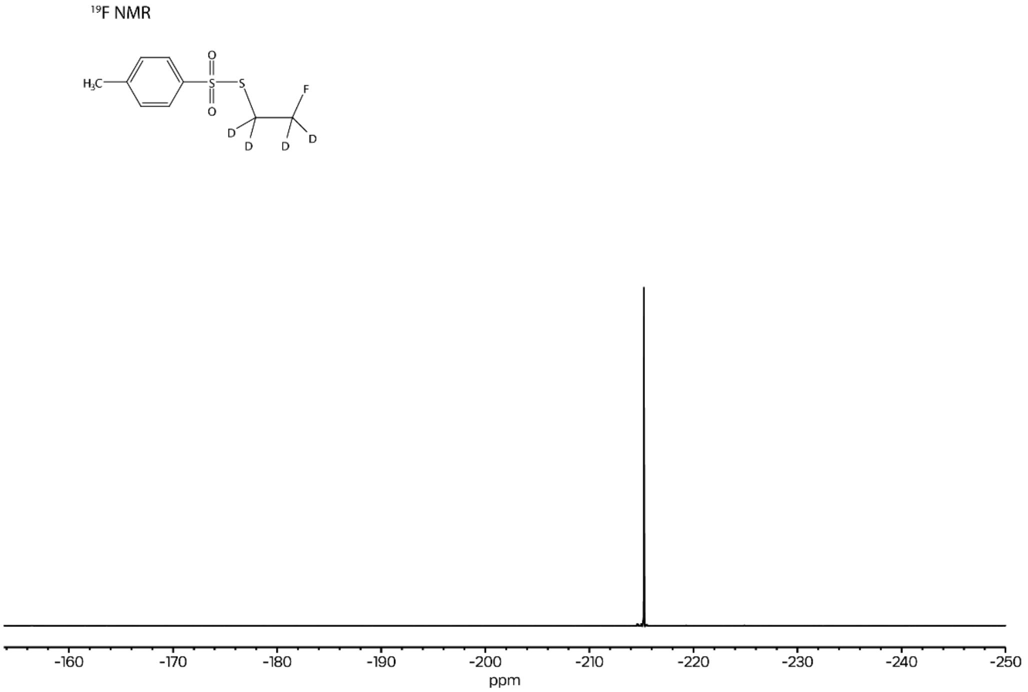


**Figure S14.** $^{1}H$ NMR spectrum of $TsSCH_2CH_2F$

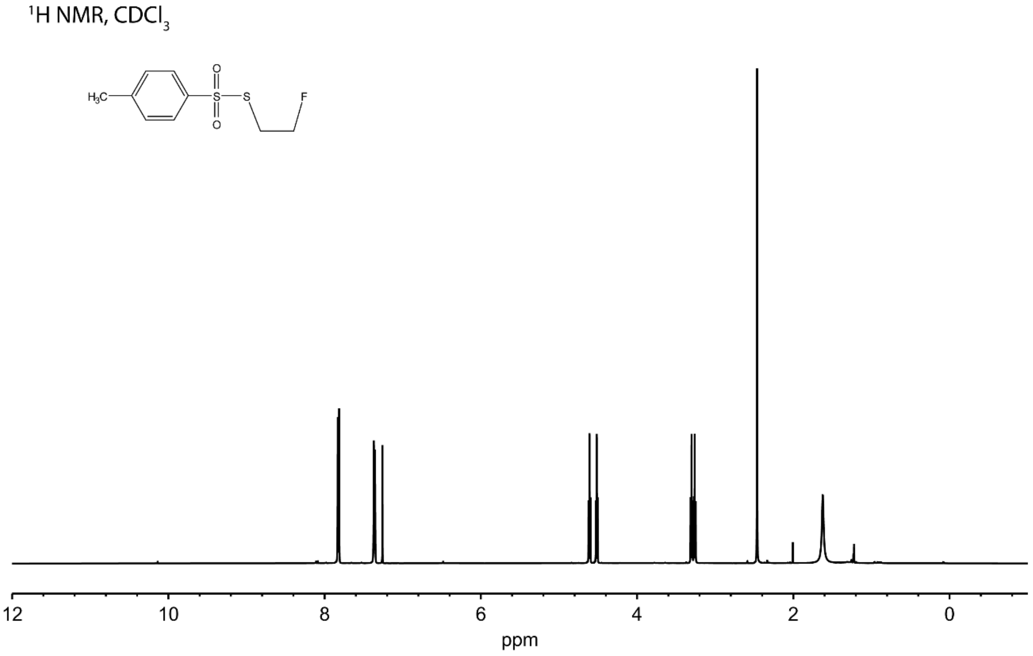

**Figure S15.** $^{13}$C NMR spectrum of $TsSCH_2CH_2F$

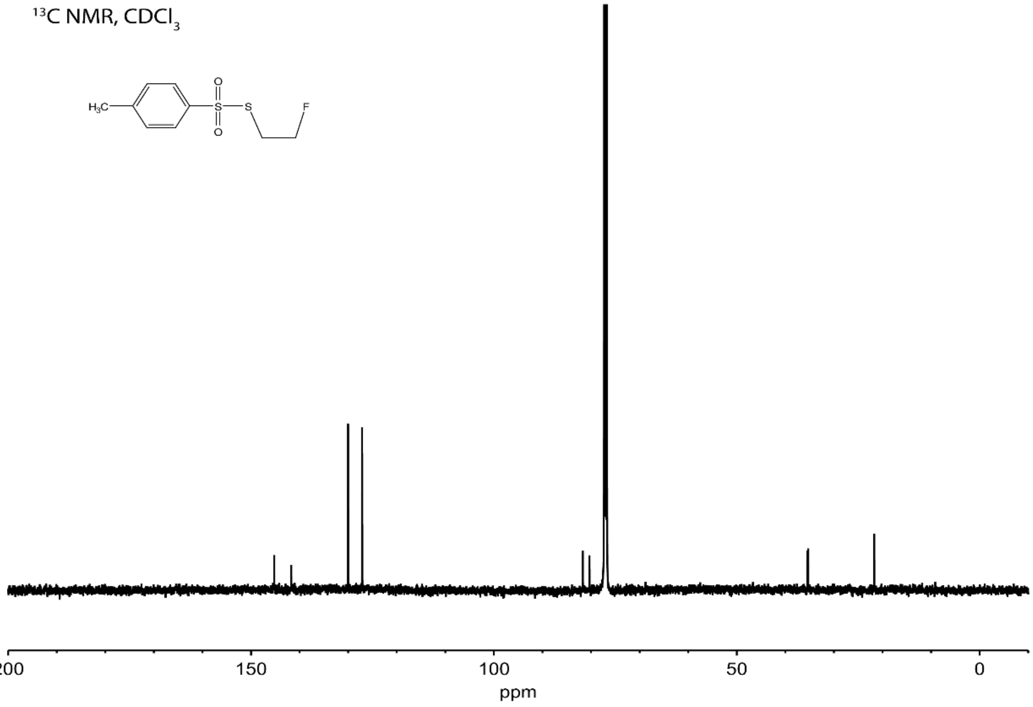


**Figure S16.** $^{19}$F NMR spectrum of $TsSCH_2CH_2F$

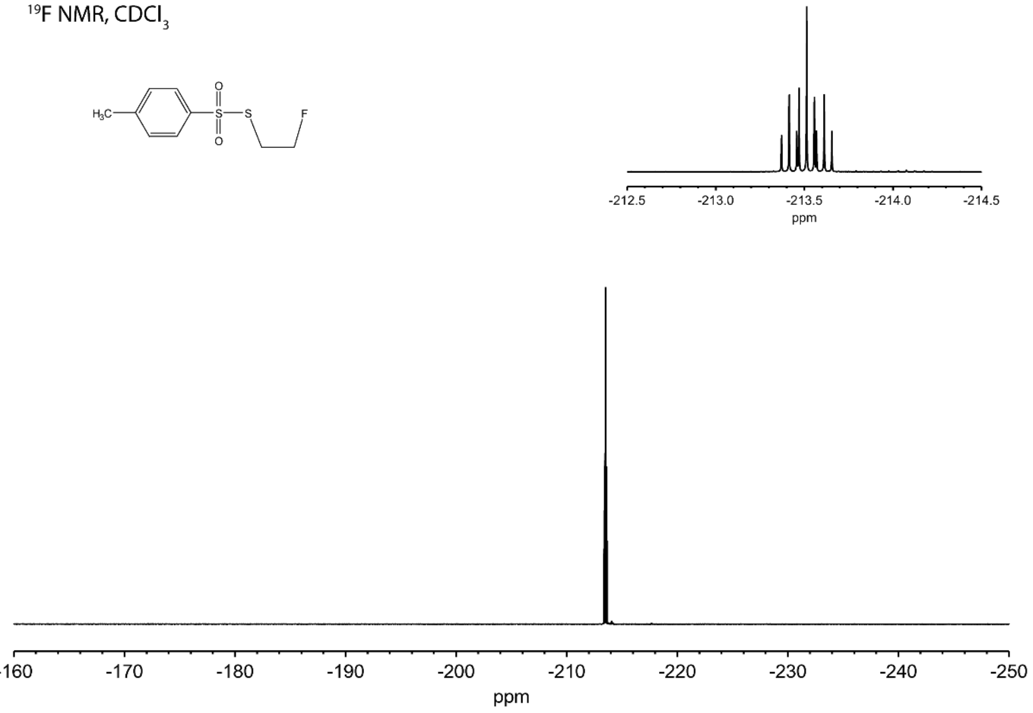
